# ECM-free patient-derived organoids preserve diverse prostate cancer lineages and uncover *in vitro*-enriched cell types

**DOI:** 10.1101/2024.10.16.618617

**Authors:** Robin Dolgos, Romuald Parmentier, Jing Wang, Raphaëlle Servant, Arnoud J. Templeton, Tobias Zellweger, Alastair D. Lamb, Kirsten D. Mertz, Svetozar Subotic, Tatjana Vlajnic, Helge Seifert, Ashkan Mortezavi, Cyrill A. Rentsch, Lukas Bubendorf, Clémentine Le Magnen

**Affiliations:** Institute of Medical Genetics and Pathology, University Hospital Basel, Switzerland; Department of Urology, University Hospital Basel, Basel, Switzerland; Department of Biomedicine, University of Basel, University Hospital Basel, Basel, Switzerland; Division of Medical Oncology, St. Claraspital, Basel, and Faculty of Medicine, University of Basel, Basel, Switzerland; Department of Urology, St. Claraspital, Basel, Switzerland; Nuffield Department of Surgical Sciences, University of Oxford, Oxford, UK; Department of Urology, Oxford University Hospitals NHS Foundation Trust, Oxford, UK; Institute of Pathology, Cantonal Hospital Baselland, Liestal, Switzerland; Clinic for Urology, Cantonal Hospital Baselland, Liestal, Switzerland

## Abstract

Patient-derived organoids (PDOs) offer new opportunities to model various cancers. However, their application in prostate cancer (PCa) has been hampered by poor success rates and overgrowth of cell types which are not representative of the patient samples. By exploiting a cohort of 164 PCa patient samples and tuning several culture parameters, we show that an extracellular matrix-free (ECM)-free culture system increases the take-rate of PDOs with luminal-like and PCa features. Single-cell RNA sequencing (scRNA-seq) reveals that ECM-free PDOs comprise cell populations associated with known PCa signatures and exhibit transcriptomic resemblance with their respective parental tumors. In addition, we define organoid-associated cell type signatures and identify markers discriminating tumors *versus* benign cells *ex vivo* and *in situ*. Furthermore, we generate the first prostate PDO single-cell atlas integrating previously-published scRNA-seq datasets and our newly-generated data. We show that Matrigel-based organoid cultures derived from primary PCa are essentially composed of benign-like epithelial cells, irrespective of the dataset or the malignant nature of the tissue of origin. In contrast, ECM-free conditions maintain heterogenous patient-specific luminal tumor cell populations and enrich in intermediate cell types. Ultimately, our work will significantly enhance the potential of PDOs in basic and translational PCa research.

## Introduction

The scarcity of effective modeling tools in prostate cancer (PCa) research poses significant challenges to understanding and treating the disease. The availability of cell lines and patient-derived xenograft (PDX) models faithfully recapitulating the complexity of PCa remains notably limited, particularly in comparison to other cancers ^1^. Illustrating this issue, the Cancer Cell Line Encyclopedia (CCLE) comprises 10 referenced PCa cell lines, representing a mere 0.52% of the total cataloged cell lines. By comparison, breast cancer, which has similar incidence rates to PCa, is represented by 97 cell lines. Moreover, the most commonly used PCa cell lines were derived from metastases, with a minority expressing the androgen receptor (AR) and being sensitive to androgen deprivation ^2^. As PCa strongly relies on AR signaling, it is essential that the models used to study the disease emulate such hallmarks ^3^. Similarly, PCa PDX model collections account for only 0.17% of all publicly- disclosed PDXs and lack certain molecular and clinical PCa subtypes ^4^.

More recently, efforts have been made to generate *ex vivo* models that better recapitulate the three-dimensional structure and heterogeneity of the tissue of origin. Notably, patient-derived organoids (PDOs) enabled significant forward strides, demonstrating enhanced functionality and molecular resemblance to the tumor of origin ^5^. To generate PDO models, cells derived from solid tumors are typically embedded in Matrigel, a murine-derived extracellular matrix (ECM) primarily composed of Laminin I and Collagen IV ^6^, and immerged in defined organ-specific medium ^5^. While PDOs generated in such conditions represent promising preclinical models in various tumor types, they are associated with limited *in vitro* tumor cell amplification, benign epithelial overgrowth, and overall low success rates in the context of PCa, thereby hindering their widespread use in translational studies ^7,8^. In spite of these challenges, a series of expandable organoid models has been successfully derived from advanced metastatic PCa specimens ^9–11^. These models, however, account for a minor subset of PCa selectively growing as successful PDOs and may represent significant outliers failing to recapitulate the broad range of PCa stages, including primary disease.

While the causes for these challenges are poorly understood, one explanation may be the multifocal nature of primary PCa, which often comprises adjacent benign glands and cancer loci ^12^. This proximity, combined with suboptimal culture conditions, may lead to the disproportionate expansion of subpopulations from an undesirable lineage, either through *in vitro* induced selection and/or plasticity mechanisms ^13^. As they may differ from their respective cell types *in situ,* the characterization of such *in vitro*-associated cellular entities remains a challenge and in-depth studies investigating these aspects are lacking. On the other hand, the high tumor cell purity of metastatic specimens, along with their enhanced aggressiveness, provides a plausible rationale for their predominance in available models. Yet this does not explain why a large majority of advanced PCa samples fail to expand in current PDO culture conditions.

The difficulties linked to generating models representing the whole clinical spectrum of PCa necessitate the research community to deviate from protocols used in other cancer types. In this study, we tackled this challenge by exploiting a cohort of 164 PCa samples and tuning several culture parameters, such as niche factors, carbon source, and the composition of the embedding extracellular matrix (ECM). We show that an ECM-free culture system enriches in PDOs with luminal-like and PCa features, while Matrigel-based organoid cultures are mainly composed of benign-like basal cells. Applying single-cell RNA sequencing, we further demonstrate that ECM-free PDOs comprise cell populations associated with known PCa signatures, exhibit transcriptomic resemblance with their respective parental tumors, and maintain patient-specific epithelial populations. Furthermore, we generate the first comprehensive prostate PDO single-cell atlas and identify organoid-associated cell type signatures, allowing refine cell type characterization *in vitro*.

## Results

### Matrigel-based culture conditions are inefficient and yield basal-like benign organoids

To generate PDO models, cells derived from solid tumors are typically embedded in Matrigel and immerged in a medium containing a cocktail of growth factors and pathways modulators ^5^. Although successful for other cancer tissues, several groups, including ours, have reported significant limitations applying such methodology to PCa samples. Following on our previous work ^7^, we persisted in our efforts and attempted to generate PDOs from an additional cohort of 136 PCa samples with diverse clinical and pathological features using previously-published conditions (**Fig.1a, b**). Half led to the short-term growth of PDOs (n=68/136 at passage 0, 50.0%); out of these, only a minor fraction (n=12/136, 8.8%) was confirmed to exhibit PCa-like features, as shown by morpho-histological characteristics (i.e. foamy-like morphology ^7^), expression of PCa-associated markers (e.g. AMACR), or genomic features (e.g. *AR* amplification) (**Fig. 1c, d**). Short-term cultures with nearly-pure tumor features originated from metastasis or transurethral resection of the prostate (TURP) specimens (**Fig. 1c, d**; **Fig. S1a**). In contrast, a minor subset of radical prostatectomy (RP)- derived organoid cultures (n=6) comprised rare tumor-like cells concomitant with numerous benign cells, highlighting challenges associated with benign overgrowth in heterogeneous primary samples.

**Figure 1.**
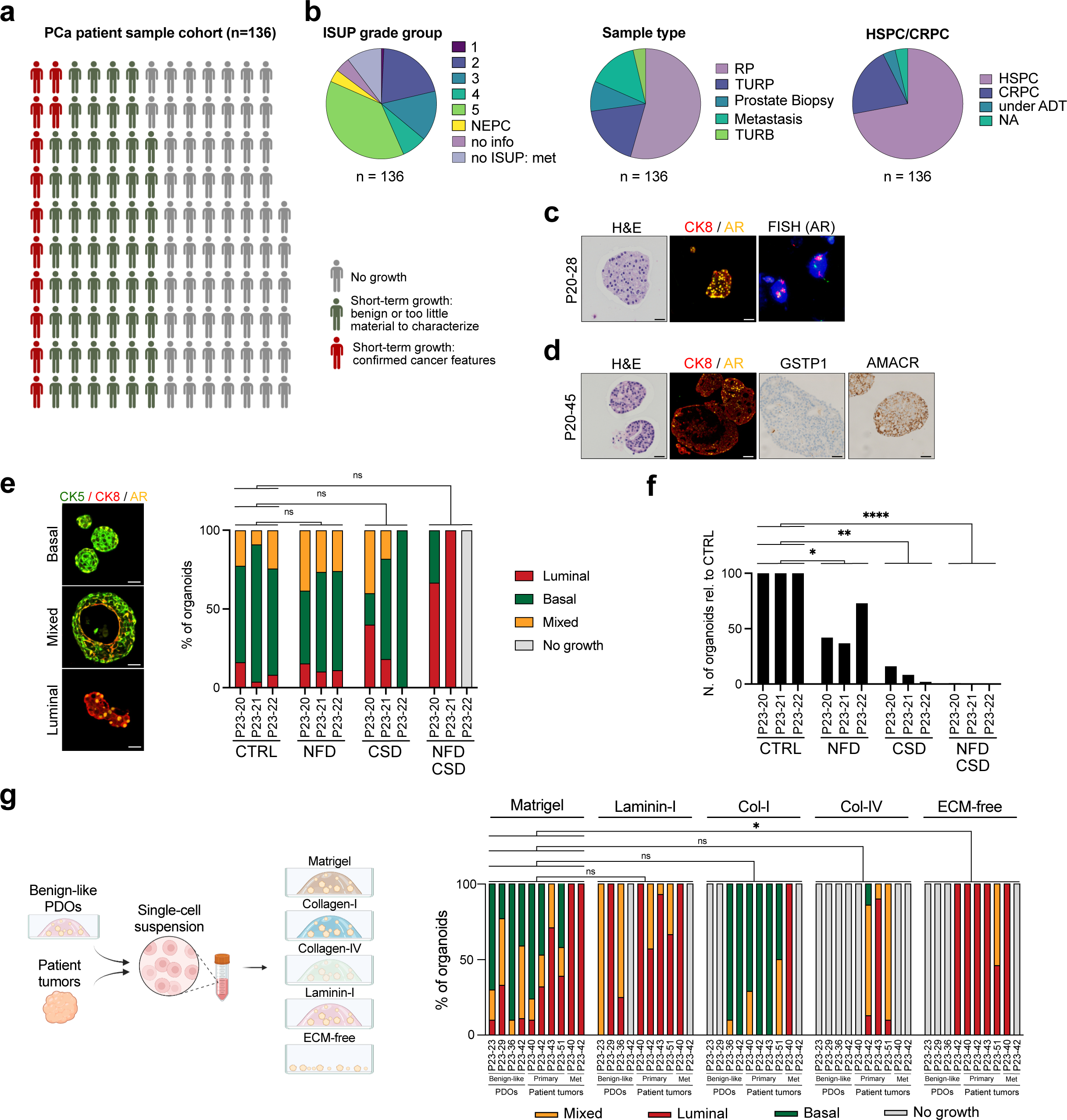
Modulating the composition of the medium and the extracellular matrix affects PDO phenotypic outcomes. **a** PCa patient sample cohort (n=136) used for PDO generation with published protocols, color-coded according to success of *in vitro* organoid culture at passage 0. Created with BioRender.com **b** ISUP grade group, sample type and castration status of PCa samples included in the study. Radical prostatectomy (RP), Transurethral resection of the prostate or the bladder (TURP or TURB). NA: not available. Hormone sensitive: HSPC, Castration-resistant PCa: CRPC. Under androgen deprivation therapy: under ADT **c - d** Examples of short-term PDO cultures with PCa features as analyzed by H&E, immunofluorescence, immunohistochemistry (IHC), and fluorescence *in situ* hybridization (FISH). Scale bars represent 50 μm. **e** Immunofluorescence-based phenotypic analysis of organoids grown in different conditions. CTRL: control medium, NFD: Niche factor deprivation, CSD: Carbon source deprivation. Organoids were categorized according to the expression of cytokeratin 5 (CK5), cytokeratin 8 (CK8), and androgen receptor (AR) as shown in left panel. Scale bars represent 50 μm. Data are represented as percentages of organoid type. Analyses were performed using a Paired t-test (*n.s*: not significant). **f** Quantification of organoids obtained from each condition, shown as relative to CTRL condition. Analyses were performed using a Paired t-test (\**p* = 0.0483, \*\**p* = 0.002, \*\*\*\**p* = < 0.0001). **g** (Left) Experimental workflow to test different extracellular matrices (created with BioRender.com). (Right) Immunofluorescence-based phenotypic analysis of organoids grown in different conditions. Analyses were performed using a Paired t-test (*n.s*: not significant, \**p* = 0.0356).

To overcome these limitations and improve culture conditions, we first sought to identify the variables responsible for benign cell expansion. Given that benign-like overgrowth is a challenge for other tumor tissue types containing adjacent tumor and benign regions ^14^, we attempted to replicate some of the approaches used to generate PDOs in these tissues. One such method involves the deprivation of niche factors leveraged to select out niche dependencies associated with normal-like phenotypes in the context of pancreatic cancer ^15^. To apply it to our context, we removed niche factors such as EGF, Noggin, and R-Spondin from the culture medium (Niche Factor Deprivation; NFD condition). In addition, as PCa cells were recently shown to switch to a lipid-based metabolism^16^, we removed carbon-sources, such as glucose, sodium pyruvate, and glutamine, from the culture medium (Carbon Source Deprivation; CSD condition). PCa progression typically associates with a loss of basal cells and an hyperproliferation of luminal tumor cells, leading to the loss of the native bi-layered epithelium and to tumors with a pure luminal phenotype ^17^; we therefore analyzed organoid phenotypic outcomes by examining the expression of basal-associated (Cytokeratin 5: CK5) and luminal-associated (Cytokeratin 8: CK8) markers in NFD and/or CSD conditions as compared to the control published conditions. Tested individually, neither condition significantly altered the phenotypic ratio and instead reduced the overall number of organoids derived from radical prostatectomy (RP) specimens, from an average of 67 to 27.6 and 7.6, respectively (n=3; Paired t test, p = 0.05, p = 0.002; **Fig. 1e, f; Fig. S1b, Table S2**). In contrast, double deprivation (CSD+NFD) appeared to limit the growth of CK5^+^ basal-like organoids (non-significant), while significantly decreasing the overall number of organoids or fully inhibiting their growth (n=3; 67 versus 1.6 organoids, Paired t test, p < 0.0001; **Fig. S1b**, **Table S2**). As the yield of luminal-like CK5^-^CK8^+^ PDOs following double deprivation was too low to consider it a success, we next sought to identify other normal-like dependencies in order to improve outcomes.

### ECM-free conditions favor the growth of organoids enriched in luminal-like and PCa features

During PCa progression, epithelial cell dynamics are accompanied by a loss of the basal lamina and by significant changes in the organization and composition of the adjacent ECM ^18^. Based on this knowledge, we reasoned that the formation and cellular composition of PCa-derived organoids might be highly dependent on the nature of the surrounding ECM.

To investigate the matter, we tested Matrigel, Laminin I, Collagen I, Collagen IV conditions, as well as complete absence of matrix (“ECM-free”: ECMf), in control published medium using 4 PDO lines previously characterized as benign-like and 6 freshly dissociated patient tumors (**Fig. 1g, left**). Although Matrigel allowed the successful generation of organoids in the largest proportion of samples (10/10), generated organoids displayed mixed phenotypes and few of them were purely luminal (**Fig. 1g; Fig S1c, d; Table S3**). Significant differences in morphology, cellular phenotypes, and viability could be observed with the other matrices tested. Collagen I strongly enriched the number of basal-like organoids generated, while only 3/10 samples grew in Collagen IV. Laminin I did not yield basal-like organoids, with mostly mixed or luminal phenotypes observed. Interestingly, Laminin-I, Collagen-IV and ECMf PDOs exhibited inverted polarity with basal cells facing inwards when present, in contrast to Matrigel-based conditions in which luminal cells faced the lumen as typically observed in the prostatic epithelium *in situ* ^17^ (**Fig. S1c**); a time-dependent reversal of normal polarity was also observed when using organoids pre-formed in Matrigel and transferred to ECMf conditions, as shown in other tissue types^19,20^ (**Fig. S1e**). Remarkably, ECMf culture did not support the growth of 3 out of the 4 benign-like PDO lines but did support the growth of purely luminal-like organoids originating from 5 out of 6 PCa patient-derived samples, highlighting the luminal selecting and/or promoting nature of these culture conditions.

To validate these initial findings, we attempted to generate PDOs in both Matrigel and ECMf conditions from 15 other PCa samples — in addition to the 6 PCa samples used in the ECM experiment — bringing the whole Matrigel/ECMf experimental cohort to a total of 21 samples with heterogenous ISUP grade group and origin (**Table S1**; **Fig. 2a**). Cell suspensions were split in two and seeded either in Matrigel or in ECMf conditions for direct comparison, whenever feasible. The growth at passage zero was of 90.5% for ECMf conditions, as compared to 76.5% in Matrigel (**Fig. 2b**). Consistent with our previous observations, organoids cultured in ECMf conditions predominantly exhibited luminal or mixed, inside-out phenotypes with AR expression restricted to CK5^-^ cells. In contrast, organoids generated in Matrigel exhibited mostly AR^-^ CK5^+^ basal-like phenotypes (**Fig. 2b, c, Fig. S2a**). Out of 17 samples attempted in both conditions, 3 grew only in ECMf conditions but failed to grow in Matrigel conditions (**Fig. 2c**). Noteworthily, all 3 samples were derived from high-grade primary samples (two RPs, 1 TURP) with minimal basal benign components, suggesting that Matrigel may restrict the growth of organoids derived from primary specimens with high tumor content.

**Figure 2.**
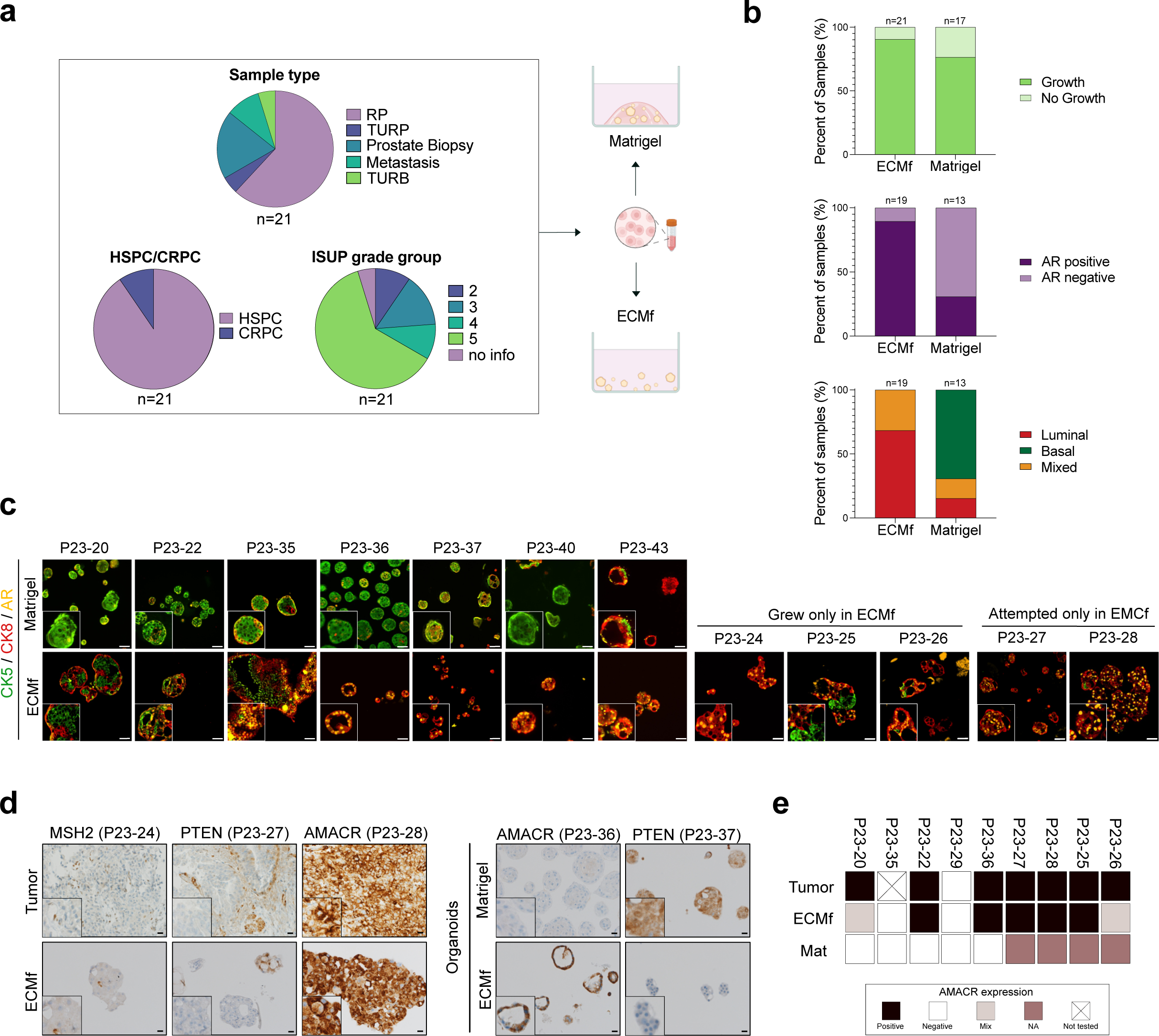
ECM-free conditions favor the growth of organoids enriched in luminal-like and PCa features. **a** Sample type, ISUP grade group, and castration status of PCa samples included in the Matrigel/ECMf comparison experiment (n=21). **b** Percentage of samples which grew, defined by the growth at passage 0 (top). AR positivity determined by immunofluorescence staining (middle). Immunofluorescence-based phenotypic analysis of organoids grown in different conditions (bottom). **c** Comparison of ECMf- and Matrigel-derived organoids derived from the same PCa samples. Organoids categorized according to the expression of cytokeratin 5 (CK5), cytokeratin 8 (CK8), and androgen receptor (AR). Insets show higher magnification of a representative area. RP: P23-20, P23-22, P23-36, P23-40, P23-25, P23-26, P23-27, P23-28; TURB: P23-24; prostate biopsy: P23-35, P23-37, P23-43. Scale bars represent 50 μm. **d** IHC analysis for the indicated antibodies. Insets show higher magnification of a representative region. Scale bars represent 20 μm. **e** Heatmap comparing AMACR status (IHC) for tumor samples and matched organoids grown in Matrigel (Mat) or ECMf conditions.

As the sole expression of luminal-like markers is not proof of the malignancy of the organoids, we next sought to further characterize them using patient-specific biomarkers identified during the pre- or post-surgery pathological evaluation of the specimens (**Fig. 2d, e; Fig. S2b**). ECMf PDOs had a striking resemblance to their respective tumors, as illustrated by the loss of the mismatch repair protein MSH2, or of the tumor suppressor PTEN, or by strong expression of the PCa biomarker AMACR ^21^. When comparing ECMf and Matrigel organoids originating from the same parental tumors, organoids with malignant features (e.g. AMACR^+^ or PTEN^-^) were only observed in the ECMf conditions (**Fig 2d, e**). Two ECMf PDO cultures, however, showed a mixed profile with cells displaying low or absent AMACR staining, suggesting that some benign cells are still maintained in these restrictive conditions (**Fig. 2e; Fig. S2b**). Finally, we confirmed that the samples which grew in ECMf but failed to grow in Matrigel displayed AMACR^+^ CK8^+^ AR^+^ phenotypes, suggesting their malignant nature. The sole Matrigel-derived PDO culture displaying a CK8^+^AR^+^ phenotype originated from a lymph node metastasis supporting that only a minor subset of metastatic specimens can grow successfully in Matrigel. Thus, ECMf conditions facilitate the growth of organoids which are enriched in luminal-like and PCa-associated features, contrarily to Matrigel.

### Single-cell RNA sequencing analysis uncovers cellular heterogeneity and tumor-like subsets in ECMf-derived organoids

Newly identified cell subtypes, referred to as hillock and club cells, have added a layer of complexity to the prostate epithelial cell lineage classification, previously restricted to basal, luminal, and neuroendocrine cells ^22,23^. Recent scRNA-seq studies have also identified these intermediate cell populations in the diseased prostate including proliferative inflammatory atrophy (PIA) and PCa ^24–27^.

To further uncover the cellular entities associated with our two culture conditions and identify putative effectors of each condition, we employed scRNA-seq on a total of 9 PDO cultures originating from 7 distinct primary PCa samples (1 Prostate biopsy, 5 RPs, 1 TURP; **Table S1**). We successfully generated libraries for matched Matrigel and ECMf PDOs for 2 out of the 7 specimens. Additionally, we generated libraries for 3 out of the 7 specimens grown only in ECMf and 2 out of the 7 specimens grown only in Matrigel. This discrepancy was due to either the failure of PDOs to grow in one culture condition or technical issues. A Uniform Manifold Approximation and Projection (UMAP) of the integrated scRNA-seq data highlighted evident segregation with residual overlap of cells according to their culture condition, driven by the presence or absence of ECM (n=39,456 cells; **Fig. 3a**). Unsupervised clustering revealed 8 distinct cell clusters, which were not driven by sample- effects or cell cycle phases (**Fig. 3b, c; Fig. S3a**). When evaluating the proportion of cells within each cluster originating from either culture conditions, we observed that clusters 1, 8, 3, and 4 were enriched in ECMf-derived cells, whereas cluster 5, 6, and 7 were enriched in Matrigel-derived cells (**Fig. 3d**). Differential expression analysis (DEA) among all clusters prompted us to classify cluster 1 and 8, which were almost exclusively ECMf derived, as “tumor-like” due to the co-expression of luminal markers (KLK3, NKX3-1) and PCa-associated markers such as AMACR, FOLH1 (encoding PSMA) and KLK4 (**Fig. 3e, Fig. S3b, Dataset 1**). In contrast, clusters 2, 5, 6 and 7 expressed both the basal marker KRT5 and the hillock marker KRT13. Finally, clusters 3 and 4 were characterized by certain club-like markers such as MMP7, CP and PSCA. To better characterize identified clusters and define their relevance to patient tumors, we assigned scores to each cluster based on previously-published PCa molecular signatures ^28–30^ (**Fig. 3f; Fig. S4a, b**). In agreement with higher expression of PCa-associated markers, clusters 1 and 8 showed significant highest scores for the three PCa signatures and for an Androgen Response signature, as compared to all other clusters (Anova test, p <2.2e-16).

**Figure 3.**
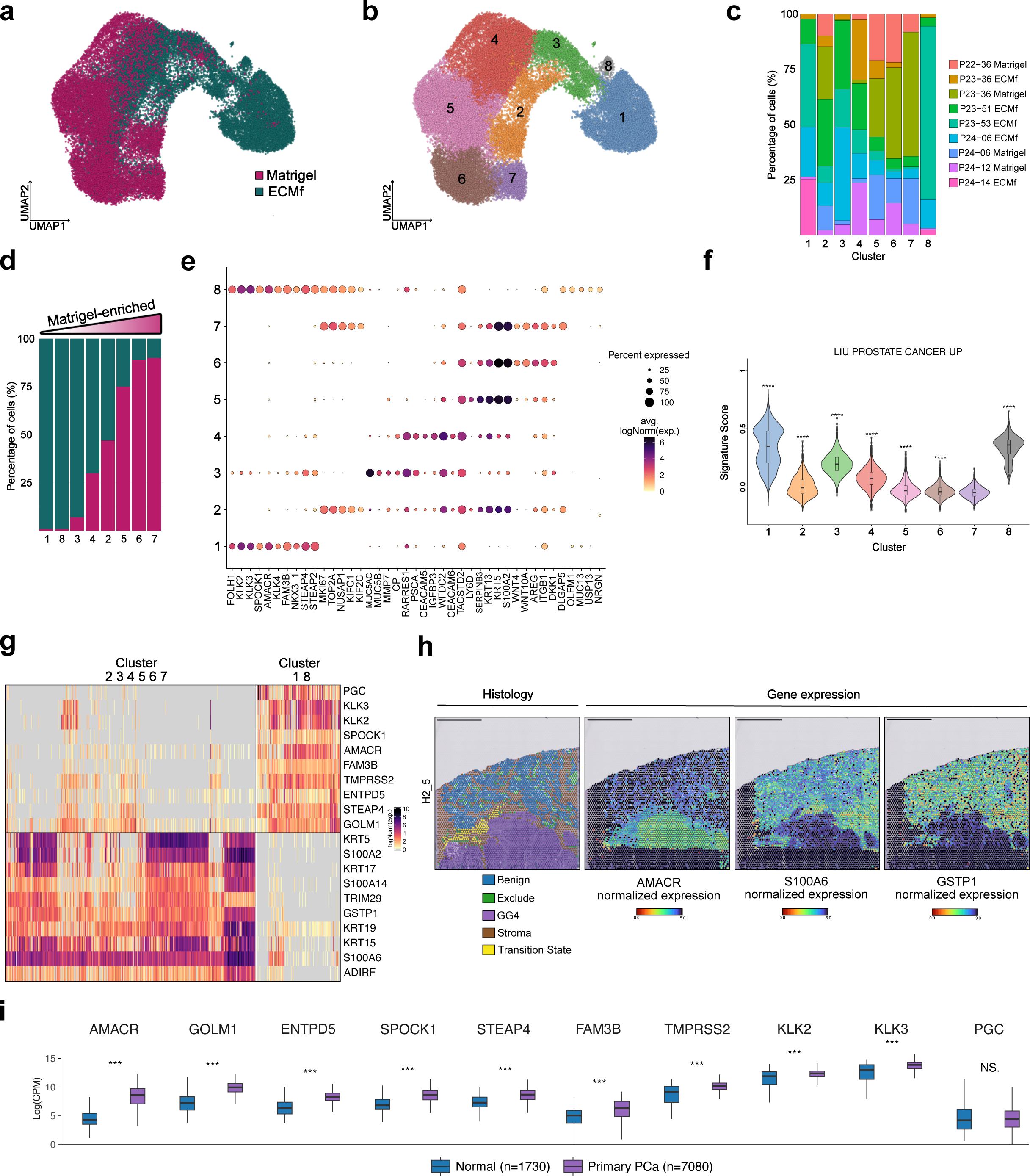
Identification of distinct benign-like and tumor-like cell populations in PCa patient-derived organoids. **a** UMAP projection of organoid-derived cells color-coded by culture condition. **b** UMAP projection of organoid-derived cells color-coded by unsupervised clustering. **c** Sample composition in each cluster. Cell counts in each cluster are normalized to 100%. **d** Percentage of cells coming from either condition per cluster. **e** Dotplot representing selected differentially expressed genes between all cell clusters. The color represents scaled average expression of marker genes in each cell type and the size indicates the proportion of cells expressing each gene. **f** Signature score of “LIU_PROSTATE_CANCER_UP” (Collection C2: Curated, gsea_msigdb) per cluster, represented as violin plots for the eight clusters (Anova statistical test, p<2.2e-16. Post-hoc test: t-test). **g** Top 10 differentially-expressed genes between clusters 2,3,4,5,6,7 and clusters 1,8, represented as a single cell heatmap plot. **h** Spatial transcriptomic (Visium ST) data from the H2_5 prostatectomy section, with spot-level pathological classification of areas (left), and log-normalized expression of AMACR, S100A6 and GSTP1. Scale bars represent 2.5 mm. **i** Expression of top 10 up genes in clusters 1, 8 in the normal prostate versus primary PCa samples using the PCa profiler bulk RNA-sequencing tool represented as box plots (Wilcoxon test; ***: P<0.001).

Interestingly, we noticed that cluster 3 also displayed moderately elevated PCa scores and AR signatures, in contrast to cluster 4 which showed similar scores as clusters 2, 5, and 7. We therefore further compared the assigned club-like clusters (4 and 3) by evaluating the expression of selected genes (**Fig. S5a**). We found that their resemblance resided in their shared expression of club-specific markers while lacking basal marker expression. Cluster 3 however, expressed low-levels of AR-pathway (KLK3, NKX3-1) and PCa-related (STEAP4, STEAP2)^31,32^ genes. Notably, a similar population was previously described as a PCa-enriched club cell population with elevated AR signaling levels ^26^. To compare their lineage trajectories, we performed Pseudotime Trajectory Analysis ^33^ using the basal/hillock clusters as starting point, as previously described ^26,22^ (**Fig. S5b**). Clusters 3 and 4 were defined by two different lineages (1 & 2), although both passed across cluster 4. Lineage 2 ended in Cluster 4 and showed terminal expression of club-specific genes such as OLFM4, suggesting a more committed club entity (**Fig. S5c**). Cluster 3 instead showed modest expression of PCa-related genes correlating with pseudotime for lineage 1 (KLK4, TMPRSS2, KLK3, FOLH1; **Fig. S5c, d**). Based on these data, we propose that cluster 3 represents a transitioning population whose transcriptomic state lies between the club and tumor lineages, while cluster 4 is composed of more committed club cells. Thus, we categorized clusters 1 and 8 as “tumor”, cluster 2, 5, 6 and 7 as “basal/hillock”, cluster 3 as “transitioning”, and cluster 4 as “club”.

Given the likely importance of integrin signaling in response to changes in the extracellular matrix, we analyzed integrin expression across distinct cellular clusters (**Fig. S6a**). Notably, unsupervised hierarchical clustering highlighted cluster-specific patterns of expression of selected integrins. Clusters 6 and 7 (basal/hillock) exhibited the highest expression of ITGB1, ITGB4, ITGA3, and ITGA6. Three other clusters (5, 2, and 4) also expressed these integrins, albeit at lower levels. In contrast, cluster 3, identified as a transitioning population, maintained most integrin expression, but showed reduced ITGB4 and ITGA6 levels. The tumor clusters had almost no expression of ITGB4, ITGA2, ITGA3 and ITGA6, maintaining only ITGB1 and ITGB5. Thus, integrins are differentially expressed in distinct ECM conditions, potentially influencing the cell types composing the resulting organoids.

### Identification of markers distinguishing between tumor and benign cells *ex vivo*

To identify biomarkers potentially allowing the clear distinction between tumors vs. non-tumor cells *in vitro* across all PDO samples, we performed DEA comparing the tumor group (1,8) against all others (2, 3, 4, 5, 6, 7) (**Fig. 3g, Dataset 1**). While confirming expression of well-known PCa-associated genes in the tumor group (e.g. AMACR, KLK3, TMPRSS2, STEAP4), this analysis did not reveal single markers highly expressed in all tumor cells and fully negative in all benign subsets. Yet GSTP1 loss of expression was observed in the vast majority of the cells present in the tumor clusters, while highly expressed in virtually all cells associated with non-tumor clusters (**Fig. 3g**). This finding is consistent with previous literature reporting that GSTP1 is frequently lost in PCa ^34^. Furthermore, DEA also revealed high and homogeneous expression of S100A6 in the basal/hillock, club, and transitioning subsets, as compared to the tumor group. To investigate whether high S100A6 expression also identified benign cells in patient tissues, we re- analyzed a spatial transcriptomic dataset generated by Erickson et al. ^35^. Consistent with our findings in PDOs, S100A6 expression was lost in AMACR^+^ GSTP1^-^ Gleason grade 4 cribriform zones, contrarily to adjacent benign/stromal areas in the same tissue sections (**Fig. 3h; Fig. S6b**). Notably, S100A6 was proposed to represent a basal cell-specific marker in a study published before the scRNA-seq era ^36^. Our data further suggest that S100A6 may represent a broader benign marker, identifying basal and intermediate (club and hillock) cells in both organoids *in vitro* and tumor samples *in situ*. To further validate the clinical relevance of the top 10 positively-enriched genes in the tumor clusters, we evaluated their expression in normal versus primary PCa samples using the PCa profiler tool ^37^. Except for PGC, all the top genes were significantly more expressed in the primary PCa cohort (n= 7080), as compared to normal prostate tissues (n=1730), further indicating their association with PCa (Wilcoxon test, p < 0.001; **Fig. 3i**).

### Organoids generated in ECM-free conditions preserve patient-specific transcriptomic features and diverse PCa lineages

To allow their implementation in precision oncology, it is crucial that PDOs faithfully emulate molecular characteristics of their respective patient tumor and do not conform to a common culture-induced entity. To address this aspect, we restricted our dataset to the tumor cluster and performed DEA between individual organoid samples, highlighting heterogeneity between samples and patient-specific transcriptomic features (**Fig. S7a, Dataset 1**). To ensure that this inter-patient heterogeneity was not *in vitro* induced, we narrowed our focus to features available for diagnostic validation at our department and identified 4 markers (ERG, ALDH1A1, VIMENTIN, MSH2) distinguishing patient-specific PDOs (**Fig. 4a**). We next investigated expression of these markers in the parental patient tumors by immunohistochemistry, enabling us to confirm MSH2 loss in the P24-14 patient sample, Vimentin overexpression in the P24-06 patient sample, ALDH1A1 overexpression in the P23-53 patient sample, and ERG^+^ tumor cells in the P23-51 patient sample, aligning with our PDO transcriptomic data (**Fig. 4b; Fig. S7b**). These data suggest that short-term PDOs cultured in ECMf conditions preserve the highly heterogeneous landscape of PCa.

**Figure 4.**
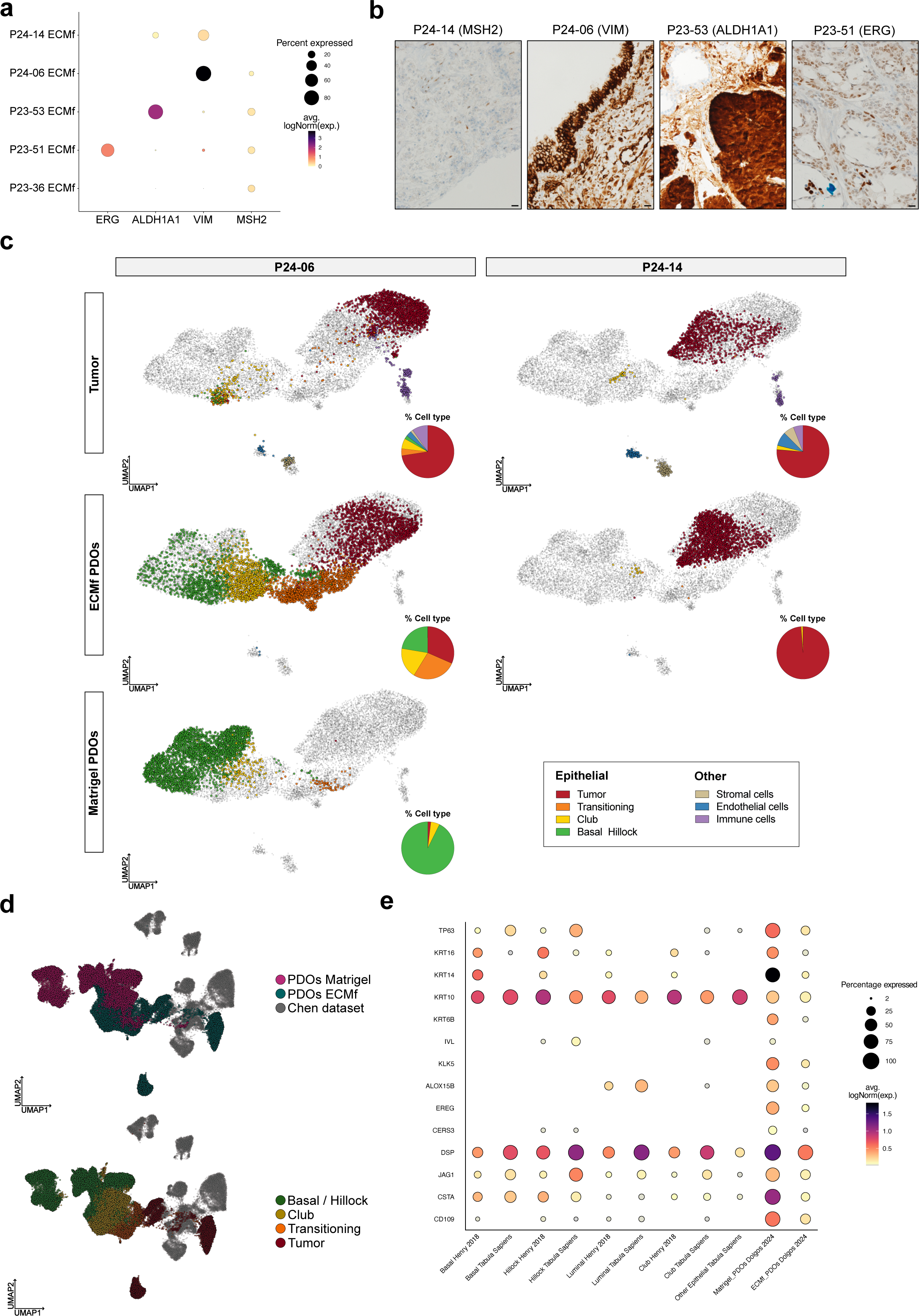
Organoids generated in ECM-free conditions preserve patient-specific transcriptomic features and diverse PCa lineages. **a** Dotplot representing expression of selected genes in clusters 1,8 (tumor) of each patient-specific ECMf organoid culture. The color represents scaled average expression of marker genes in each cell type and the size indicates the proportion of cells expressing each gene. **b** IHC analysis for the indicated antibodies in corresponding patient samples. Scale bars represent 20 μm. **c** UMAP projection of organoid-derived cells (ECMf or Matrigel) and matched patient samples separated by patient. Each UMAP is accompanied by pie charts depicting the proportion of each cell type per sample and condition. **d** UMAP projection of generated organoid data integrated with the Chen dataset. Top: Cells are color-coded by dataset. Bottom: Cells are color-coded by assigned cell-type signatures using the singleR classifier. **e** Dotplot representing expression of selected genes associated with the “GOBP Keratinocyte Differentiation” signature (Collection C5: Ontology, gsea_msigdb). The color represents scaled average expression of marker genes in each cell type and the size indicates the proportion of cells expressing each gene. Genes expressed by less than 2% of the cell population are not displayed.

To further investigate the similarities and differences between matching parental tumor and PDOs, we sequenced both the original tumors and the corresponding organoids for two of the seven PCa specimens (**Fig. 4c**). Notably, P24-06 tumor (treatment-naïve RP, ISUP4) formed organoids in both ECMf and Matrigel conditions, while P24-14 (castration-resistant TURP, ISUP5) only grew in ECMf conditions. After data integration, separating the samples by patient revealed a major overlap of the P24-06 tumor with its ECMf counterpart in terms of tumor cells and an important proportion of club, transitioning, and basal/hillock cell types in ECMf-organoids. Notably, these intermediate populations were present in minor proportion in the parental sample, suggesting an *in vitro*-specific enrichment. In contrast, Matrigel-derived organoids only minorly overlapped with the benign cell populations of the parental tumor, clustering on their own and assigned as basal/hillock cells, which were a minority in the parental tissue. In the case of P24-14, we observed a nearly perfect overlap between the patient tumor and ECMf-derived organoids, in terms of both cancer and club-like populations. For both patients, besides rare endothelial cells, non-epithelial populations (i.e. immune, mesenchymal) were absent in organoids, highlighting the epithelial-selective nature of these culture conditions. To expand the relevance of our findings to larger and independent cohorts of patients, we integrated our PDO samples amongst epithelial scRNA-seq profiles of 13 PCa tumors generated by Chen and colleagues ^25^. After assigning cell types to the Chen dataset by cell-type annotations (**Fig. S7c**), we observed that ECMf PDOs partially overlapped with both luminal and basal cells of PCa tissues *in situ* (**Fig. 4d**, top panel). On the other hand, Matrigel PDOs clustered almost entirely by themselves or aside basal cells, suggesting enrichment of basal-like cell populations and *in vitro* specific features (**Fig. 4d**). Leveraging our previously-generated signatures, we confirmed that ECMf PDO cells clustering near or with Chen luminal cells were classified as “tumor”, while Matrigel PDO cells clustered almost entirely on their own and were classified as “basal/hillock” (**Fig. 4d**, bottom panel).

Taken together, these data suggest that ECMf-derived organoids comprise basal/hillock, club, transitioning and tumor cell subsets highly resemblant to their original patient tumor and representative of diverse PCa lineages. In contrast, Matrigel fails to maintain tumor-like cells and produces *in vitro* basal-like transcriptomic profiles that are highly divergent from PCa patient samples.

### Matrigel induces keratinocyte-like lineage programs

Given that cells derived from the Matrigel condition mostly clustered far apart from PCa samples in the Chen dataset as well as in our cohort, we next aimed at characterizing these divergent transcriptomic features. As previously described by our group and others ^7^, a large subset of Matrigel-derived organoids exhibits a squamous-like histo-morphology which can be accompanied by keratinization features (**Fig. S7d**). Consistent with these observations, basal/hillock cell types which are largely enriched in Matrigel conditions, displayed a high keratinocyte differentiation signature score (**Fig. S7e**). To understand whether these features were representative of the normal prostate, we compared the most relevant genes from the keratinocyte differentiation signature across all epithelial cell types comprised in two normal human prostate datasets ^22,38^, along with our Matrigel and ECMf data (**Fig.4e**). These analyses highlighted the aberrant expression of numerous keratinocyte-associated genes (e.g. CSTA, KLK5, EREG) in Matrigel PDOs, which were absent or expressed at much lower levels in normal prostate datasets and in ECMf organoids. Thus, Matrigel-derived PCa organoids exhibit transcriptomic features that are not representative of the normal prostate epithelium, suggesting potential *in vitro*-induced transcriptomic features.

### An integrative single-cell atlas of prostate PDOs further highlights the limitations associated with matrigel-based organoid culture

To broaden our comparison between organoid models, we integrated our newly-generated dataset with all identified publicly-available scRNAseq data of prostate PDOs. Specifically, we included three single-cell datasets from Matrigel-based organoids derived from normal prostate ^39^, PCa samples ^24^, and matched normal and PCa samples ^26^ (n= 31 organoid samples total; n=96,038 cells). Additionally, we generated scRNA-seq profiles of two PCa organoid models previously generated in our laboratory (P20-11 and P20-23 PDO xenograft organoids, see *Methods*) as well as the MSK-PCa2 organoid line ^9^. All three models were cultured in both ECMf and Matrigel conditions and further referenced as long-term lines in this manuscript. The general UMAP representation of the integrated data revealed three major groups: one consisting solely of cells from the ECMf condition, another exclusively containing Matrigel-derived cells, and a third with cells from both conditions (**Fig 5a, left panel).** Interestingly, long term lines, which were initially generated and maintained in Matrigel, clustered together regardless of the culture conditions (**Fig 5a**, right panel). This may indicate that, at least at this timepoint, tumor cells that have already adapted to *in vitro*/*in vivo* environments (i.e. xenograft and/or Matrigel-based organoid culture) do not undergo transcriptomic shifts and/or selection of certain cell subsets when transferred to ECMf culture conditions. When comparing the malignancy of the tissue of origin used to generate the PDOs, we observed that matched tumor/normal samples from the Song dataset overlapped almost perfectly, further indicating that Matrigel favors the growth of benign cells regardless of the presence of tumor cells in the clinical specimen (**Fig 5b**).

**Figure 5.**
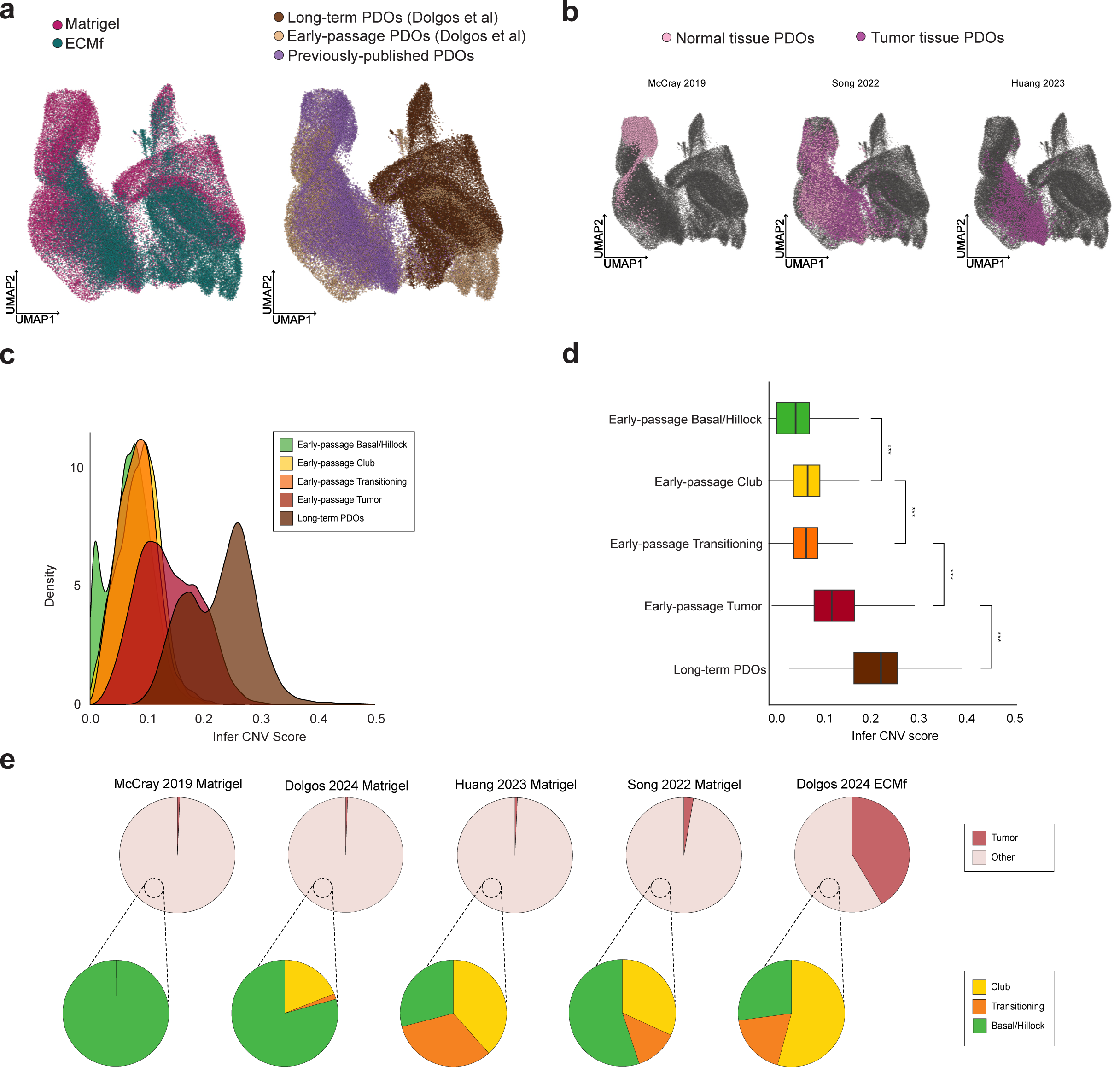
An integrated scRNA-seq atlas of organoids derived from normal and PCa patient samples. **a** Integrated UMAP projection of organoid-derived cells associated with this study (Dolgos et al: early passage and long-term PDOs) and previously published studies including the McCray, Song, and Huang datasets. UMAPs are color-coded by culture condition (left) and data origin (right). **b** UMAP projection separated by dataset and color-coded by source (normal or tumor) for PDO generation. **c** Density plot representing infer CNV scores per cell type regrouping all datasets. **d** Box plots representing infer CNV score per cell type. Wilcoxon test; ***: P<0.001. **e** (top) Pie charts of individual datasets color-coded by percentages of tumor cells and other non-tumor cell types. (bottom) Pie charts depicting the cellular composition of other cell types for each dataset.

To further validate that our signatures reliably discriminate between tumor, and benign cells *in vitro*, we inferred copy number alterations using InferCNV (**Fig. 5c, d**). Additionally, we aimed to determine whether transitioning cells represent potential tumor cells undergoing transcriptomic shifts to adapt to *ex vivo* culture. We observed significantly higher CNV scores in cells classified as tumor by our signature, which were almost entirely derived from ECMf conditions (p < 0.001, Wilcoxon test). In comparison, basal/hillock, club, and transitioning all showed lower CNV scores. Thus, transitioning cells do not appear to be genomically engaged as tumor cells, although they express certain PCa-associated markers. Applying our previously defined *in vitro* signatures on this integrated atlas allowed us to account for the proportion of cell-types in each dataset within the atlas (**Fig. 5e).** Strikingly, only ECMf short-term PDOs comprised a significant proportion of tumor cells. Cells not classified as tumor were essentially classified as basal/hillock in Matrigel PDOs, while club and transitioning cells were present in heterogenous proportions depending on the dataset. In particular, organoids from the Huang dataset comprised high proportions of cells with a club and transitioning phenotype, which may be explained by their use of tissues affected by PIA enriched in such cell types ^24^.

Collectively, these data further show that ECMf conditions favor the maintenance of tumor, club and transitioning cell populations, while Matrigel conditions largely favor the expansion of organoids with an aberrant basal/hillock phenotype irrespective of the tissue or origin.

## Discussion

Matrigel-based culture systems have allowed the development of PDO models faithfully recapitulating various human tissues and cancer types. Low yield of cells post-digestion, coupled with low tumor cell amplification and benign overgrowth *in vitro*, have however limited their application in the context of PCa. Taking these striking differences with other tissues into consideration, we propose an ECM-free system increasing PCa PDO success rates, maintaining lineage specificities and limiting benign overgrowth (see summary of findings in **Fig. 6**).

**Figure 6.**
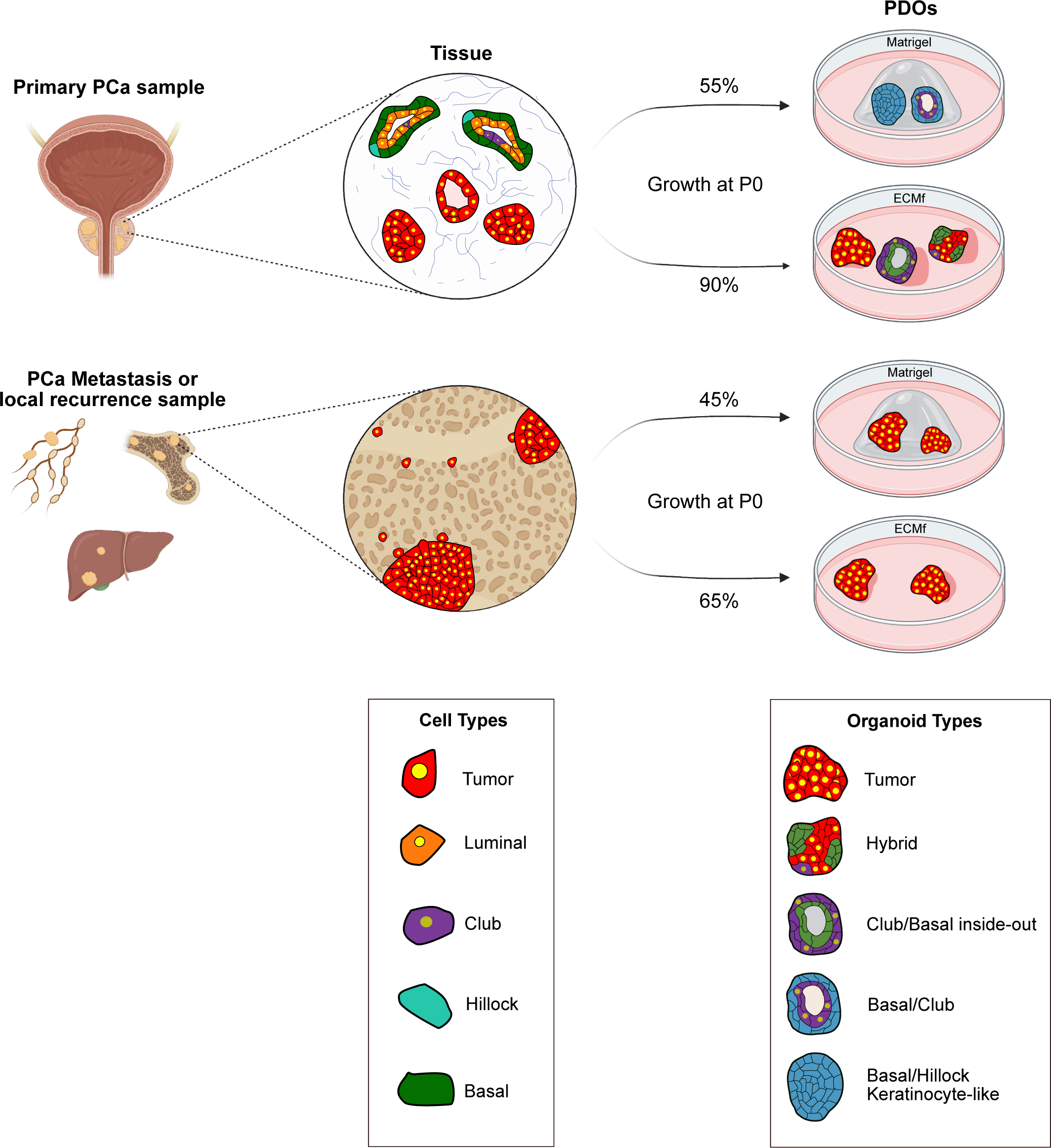
Summary of findings: anticipated outcomes of PCa PDO generation in Matrigel and ECMf conditions. Graphical overview of outcomes in terms of growth at passage 0 and cellular composition of organoids derived from primary samples (top) or metastasis specimens (bottom). “Growth at passage 0” percentages are inferred based on cohorts and analyses associated with this study and represent putative approximative values. Highlighted nuclei refer to AR+ cells and intensity of color indicate putative levels of expression. For model simplicity, non-epithelial cells (e.g. stromal, immune, endothelial) are not represented. Created in part with BioRender.com.

Several groups have previously used ECM-free systems for organoid culture and drug-testing endeavors ^9,40^. Such systems are typically favored for experimental simplicity as they facilitate seeding of cells for high throughput drug screening and avoid the use of costly and non-standardized hydrogels such as Matrigel. Investigators have also shown that ECM composition and stiffness may alter drug treatment response ^41,42^. These studies, however, did not address the possible selection of certain cell types and/or phenotypic shifts associated with specific culture conditions. Bridging this gap, our study clearly demonstrates the maintenance of patient-specific PCa cell populations in defined organoid culture conditions lacking serum and Matrigel. Moreover, we show that extensive characterization is necessary due to the enrichment of certain non-cancerous cell types in such conditions. In this framework, we define transcriptomic signatures discriminating tumor cells and other epithelial cell types *in vitro*, thereby allowing robust characterization of PDO cultures.

Beyond their inefficacy, our study clearly demonstrates that Matrigel-based culture conditions can be detrimental to prostate tumor cell survival *in vitro* and induce aberrant lineage programs in seemingly normal prostate cells. Illustrating so, we show that short-term PDO cultures, generated in such conditions by our group and others, do not comprise significant tumor populations and are enriched in benign-like cell populations which are not representative of prostate tissues *in vivo*. While the presence of *in vitro*-associated cells was already suggested in previous studies ^24,26^, we show that cells classified as “basal/hillock” in Matrigel conditions exhibit aberrant transcriptomic features resembling that of keratinocytes which are highly divergent from basal or hillock cells previously described in normal prostate tissues. Noteworthily, the challenges associated with maintenance of tumor cells in such conditions might be explained by the inherent nature of Matrigel, a basement membrane-like extract which isolates the cells from one-another. In the normal prostate, basal cells produce and are in direct contact with the basement membrane, while luminal cells lie on top of the basal layer ^17^. In contrast, PCa progression associates with the loss of the basal cell layer and degradation of certain basement membrane components. Based on this background and our data, we may speculate that most tumor cells are unable to survive and thrive in Matrigel (or other matrices with similar biological effects) due to a lack of cell-to-cell contact and forced contact with basement membrane components. In contrast, a minority of PCa samples may possess specific molecular traits and an advantage of growth in such conditions.

Relative to our findings, we may also hypothesize that benign epithelial cell survival is tightly linked to epithelium polarity, which may limit their survival and expansion in ECMf conditions lacking cell-ECM integrin binding. Tumor cells, in contrast, may acquire independence towards polarity during carcinogenesis, as supported by the loss of Integrin Beta 4 (ITGB4) observed in PCa tissues and in our PDO tumor cell populations ^43^. Moreover, the vast majority of benign cells surviving in ECMf conditions are club-like cell populations; these cells may possess inherent traits allowing them to resist in stress-inducing environments, as previously shown in the lung ^44^. Importantly, club cells have been proposed to represent precursors of luminal cells and express markers typically associated with a luminal phenotype, such as KRT8 ^22^. While these features may lead to the misclassification of these cells, most club cells associate with low AR response signatures which distinguish them from luminal (tumor) cells. The enrichment of these cell populations *in vitro* may therefore represent a significant limitation to test the efficacity of AR-targeted compounds in organoid cultures generated from heterogenous PCa samples. On the other hand, it provides a unique opportunity to study the role and functional properties of these cells *in vitro*. Finally, it is worth highlighting that PCa cells may be difficult to distinguish from healthy committed luminal cells in scRNA-seq data. While we cannot fully exclude the presence of a subset of healthy luminal cells in our assigned tumor cluster, we provide a combination of evidences (i.e. expression of PCa markers, similarity to PCa tissues, higher inferCNV profiles) clearly pointing to the maintenance of tumor cells in our culture system. To the best of our knowledge, our work is the first study demonstrating such at the single cell transcriptomic level, allowing an in-depth dive into *in vitro*-associated cellular heterogeneity and cell dynamics.

We acknowledge that ECMf PDOs go against certain aspects of traditional organoid definitions, where ECM-isolated single cells with multipotent capacities give rise to multicellular 3D structures. However, PCa tissues have lost normal epithelium regeneration capabilities and are instead formed by clonal amplification of malignant cells with more restricted cell fates. Thus, we ultimately aim at providing favorable conditions for tumor cells to thrive *ex vivo*, while maintaining cell types which closely resemble their respective tumor. In this framework, we show that ECM-free culture allows the generation of organoids derived from diverse types of PCa samples, regardless of their stage or molecular features. Although this culture system increases the yield of tumor cells representative of patient tumors, their limited amplification remains a challenge. While short-term experiments are feasible (up to five passages), we have not been able to amplify the lines enough to store them and share with other groups. Finally, success-rates for organoids derived from PCa metastases must be increased to enable their broader application in personalized medicine. Thus, future research efforts should focus on refining medium formulations as well as tuning biological and physical parameters to further improve fidelity and longevity of PCa PDO models.

## Methods

### Patients and clinical samples

All PCa samples were obtained under an approval by the Ethics Committee of Northwestern and Central Switzerland (EKBB/EKNZ 37/13) from patients operated at the University Hospital of Basel (USB) the St. Claraspital Basel (SCB), and Kantonsspital Baselland (KSBL), Switzerland. Patient candidates for tissue collection were identified at the multidisciplinary tumor conferences prior to tumor resection based on clinical, radiological and pathological features. A total of 164 samples were obtained from 162 patients and were enrolled in organoid experiments. Detailed clinical and pathological characteristics of the patients and associated samples are described in **Supplementary Table S1.**

### Tissue processing for organoid culture

Upon reception, tumor specimens were mechanically triturated with scissors in adDMEM/F- 12 (ThermoFisher, Cat: 12634010). Once fragments smaller than 1 mm³ were obtained, the mixture was transferred to a 15 mL tube, centrifuged at 350 G for 5 min, and resuspended in 10 mL of enzymatic digestion mix (1 mg/mL Dispase, 2 mg/mL Collagenase IV, 10 mg/mL DNase I, and 1 mM ROCK inhibitor). The content was transferred to a 75 mm Petri dish and placed on a rocking platform at 37 °C (20 rpm). Every 15 min, the mixture was pipetted up and down with a wide-bore serological pipette. This digestion step was allowed to last between 10-60 min, with stromal-rich samples taking significantly longer to release cells compared to epithelial-rich samples. The mixture was then passed through a 100 µm filter and transferred to a 50 mL tube, where 20 mL of adDMEM/F-12 was added. Cells were centrifuged at 350 G for 5 min, and finally, resuspended in 1 mL of adMEM/F-12.

### Organoid culture

For Matrigel-based culture, patient samples were processed and embedded in Matrigel following previously-published protocols ^7^. For ECM-free culture, patient samples were processed the same way and cells were seeded at 20,000 cells/cm^2^ in ultra-low adherence plates (Corning Costar). The organoid medium was identical to the previously published composition ^7^, and was changed every 3 days, supplemented with either 1 nM DHT or 0.1 nM DHT for hormone-sensitive PCa (HSPC) or castration-resistant PCa (CRPC) samples, respectively. Organoid cultures were passaged after 7-14 days of culture, depending on doubling time.

### Organoid passaging

Matrigel-derived organoids were passaged as previously described ^7^ .To isolate single cells from organoids grown in ECM-free conditions, the entire contents of the wells were transferred to a 15 mL tube, adjusted to a final volume of 10 mL with DMEM/F-12, and briefly centrifuged at 270G to separate single cells that have not formed organoids from multicellular 3D structures. The supernatant, containing most of the single cells, was carefully removed without disturbing the organoid pellet, leaving approximately 1 mL of residual volume. The total volume was then adjusted to 10mL with adDMEM/F-12, organoids were spun down at 350 G for 5 min and the supernatant was discarded. Organoids were further digested with 2 mL of TrypLE 1X. The digestion was considered complete when more than 80% of the organoids were reduced to single cells and should not exceed 10 min.

### Extracellular Matrix testing

Matrices were used according to manufacturer recommendations. Matrices used in this study were: Collagen-IV (Biotechne 3410-010-02), Laminin-I (Biotechne 3446-005-01), Collagen-I (Biotechne 3447-020-01). Cells were re-suspended in matrices at 750 cells/µl and the medium was changed at day 4. Viability and phenotypic analysis was performed at day 7 (see below).

### Viability analysis

Cells were seeded either in Matrigel or in ECMf conditions at 2500 cells per well in ultra-low adherence 384 well plates. Cell viability was assessed using the Cell-Titer-Glo3D luminescence assay (Promega G9681) according to the manufacturer guidelines. All viability treatment experiments were conducted in triplicates.

### Histology, immunohistochemistry, and immunofluorescence

Embedding of cultures with low numbers of organoids was performed as described in Fuji et al ^45^. Histological analysis of the samples was performed with standard hematoxylin and eosin (H&E) staining. Immunohistochemistry (IHC) and Immunofluorescence (IF) stainings were performed as previously described in Servant et al ^7^. Immunofluorescence images were captured using a Nikon Ti2 microscope (DBM Imaging Facility). For whole-mount immunofluorescence, cells were transferred to ultra-low adherence 384 well plates in PBS. A fixation step was performed using 4% Formalin for 1 hour at 37°C. After washing, cells were permeabilized with 0.2% Triton-X (Bio-Rad #1610407 ) for 20 minutes at RT. Normal Goat Serum (Vector Laboratories S1000) at 5% was incubated for 1h for blocking. The primary antibody mix was added and incubated overnight at 4°C. The secondary antibodies and DAPI were added the next day and incubated for 2 hours at RT. The plates were imaged using the Nikon Crest-X-Light V3. Details of all antibodies and dilutions are provided in **Table S4.**

### Determination of growth and characterization of tumor-associated features in the matrigel cohort

For the Matrigel cohort (Fig. 1a-d), criteria of success and/or failure were determined according to the following rules. “No growth” was determined when no organoid formation was observed by Brightfield microscopy at passage 0. “Short-term growth: benign or too little material” was determined when PDOs were visible at passage 0 but histological analysis was technically impossible or when PDOs exhibited a clear squamous morphology as described in Servant et al ^7^. “Short-term growth: confirmed cancer features” was determined when over 10% of PDOs exhibited cancer-like morphologies with PCa features demonstrated by either a “foamy-like” morphology, fluorescence *in situ* hybridization (FISH), or IHC staining for PCa associated markers. *AR* amplification was detected by FISH using the Vysis LSI Androgen Receptor Gene (Xq12) SpectrumOrange and the CEPX SpectrumAqua probes (Abbott).

### Imaging and quantification of organoids number and phenotype

The total organoid number was counted manually directly in 384 well-plate wells, in triplicates per condition (Fig. 1f). The organoid phenotype was determined either using whole-mount confocal immunofluorescence (fig. 1f, g) or immunofluorescence of formalin-fixed organoids embedded in paraffin blocks (Fig. 1g; Fig. 2b). All quantification data are reported in **Table S2** and **S3**.

### Long-term organoid models

The MSK-PCa2 organoid line was kindly gifted by Dr. Yu Chen and initially generated from a CRPC bone metastasis sample ^9^. P20-11 and P20-23 PDOs were established from a lung metastasis specimen (HSPC patient) and a TURP specimen (CRPC patient), respectively and initially reported in ^7^. PDO-xenografts (PDOXs) were then obtained by subcutaneous injection of dissociated PDO cells in NSG male mice, with or without testosterone pellet, respectively. PDOX tumors were then dissociated and cultured as organoids (PDOXOs).

### Processing of single-cell suspensions for scRNA-seq analysis

Following sample digestion into single-cell suspensions, cell count and viability was assessed using Trypan Blue staining with automated cell counting. For samples with viability below 60%, an Annexin-V-based magnetic sorting assay (Miltenyi Biotech, 130-090-101) was used to enrich for live cells. After this step, samples with viability below 80% or with fewer than 50,000 viable cells were considered not suitable for scRNA-seq experiments. As 10X genomics protocols for scRNA-seq require clumps to be absent from the loaded cell suspension, a filtering step using a pre-separation filter of 30µm (Milenyi Biotec, Cat: 130- 101-812) was used, whenever needed. For this purpose, cells were first transferred to a 15mL tube, resuspended in a total volume of 5mL of adDMEM/F-12 and passed through the filter. Finally, cells were spun down at 350 G for 5 min, the supernatant was discarded and cells were resuspended in 1mL of the same media and counted again with Trypan Blue.

### Library preparation and alignment of single-cell fixed RNA-Profiling data

Sequencing libraries were prepared according to the Chromium Fixed RNA Profiling Multiplexed Samples User Guide (10x Genomics, Revision D). Briefly, samples meeting viability and cell number criteria were fixed at 4°C for 20 hours and stored at -80°C, following the manufacturer’s guidelines (Fixation of cells & nuclei for Chromium Fixed RNA Profiling). Batches of four samples were multiplexed using the Human Transcriptome 4 probe set. Following probe hybridization and washing, fixed cells were counted using Trypan Blue for each sample within a batch and then pooled in equal numbers to achieve a concentration of 1,750–2,500 cells/µL. Each batch of four samples was loaded into a Chromium Microfluidic Chip to target a recovery of 40,000 cells (10,000 cells per sample). Immediately after Gel Beads-in-Emulsion (GEM) generation, ligation and extension of captured mRNA were performed to incorporate Unique Molecular Identifiers (UMIs), 10x GEM Barcodes, and partial Read 1T sequences. Pre-amplification of ligated products was performed after breaking the GEMs and SPRIselect beads (Beckman Coulter, B23317) were used to select for pre-amplified products. Finally, libraries were constructed by adding Illumina P5/P7 adapters and i5/i7 sample indexes, along with the Illumina TruSeq Read 1 (Read 1T) and Small Read 2 (Read 2S) sequences, via Sample Index PCR (8 cycles).

### Library preparation and alignment of single-cell 3’ gene expression data

Single-cell gene expression libraries were prepared according to the Chromium Next GEM Single Cell 3’ v3.1 Dual Index User Guide (10x Genomics, Revision B). Briefly, samples meeting viability and cell number criteria were resuspended at a concentration of 700–1,200 cells/µL and then loaded into the Chromium Next GEM Chip G to reach a recovery of 10,000 cells per sample. Immediately following GEM generation, reverse transcription of captured mRNA was performed to incorporate barcoded primers, including Unique Molecular Identifiers (UMIs) and 10x GEM Barcodes to the captured mRNA. Following Post-GEM-RT cleanup, PCR amplification of the first-strand cDNA was performed by PCR (11 cycles), followed by SPRIselect bead-based size selection (Beckman Coulter, B23317) to enrich for cDNA fragments of the desired size. Enzymatic fragmentation, end repair, A-tailing and ligation steps were performed immediately after. Finally, libraries were constructed by adding Illumina P5/P7 adapters and i5/i7 sample indexes via Sample Index PCR (9-10 cycles).

### Sequencing

All generated libraries were evaluated for quantity and quality using a Thermo Fisher Scientific Qubit 4.0 fluorometer with the Qubit dsDNA HS Assay Kit (Thermo Fisher Scientific, Q32851) and an Advanced Analytical Fragment Analyzer System using a Fragment Analyzer NGS Fragment Kit (Agilent, DNF-473), respectively. The libraries were pooled with other 10 x Genomics libraries and sequenced with a loading concentration of 300 pM, symmetrical paired-end and dual indexed, using various illumina NovaSeq 6000 Reagent Kits v1.5 (100 or 200 cycles), on an illumina NovaSeq 6000 instrument. The read set-up was as follows: read 1: 29 cycles, i7 index: 10 cycles, i5: 10 cycles and read 2: 89- 91cycles. The quality of the sequencing runs was assessed using illumina Sequencing Analysis Viewer (illumina version 2.4.7) and all base call files were demultiplexed and converted into FASTQ files using illumina bcl2fastq conversion software v2.20. All steps post library preparation were performed at the Next Generation Sequencing Platform, University of Bern.

### Alignment and demultiplexing of single-cell fixed RNA-Profiling data

Genome alignment, UMI counting and sample demultiplexing of the sequenced data for each batch of four samples was conducted using *cellranger multi pipeline* (v7.2.0) with default parameters, according to user guidelines available on the manufacturer’s website (www.10xgenomics.com/support/software/cell-ranger/latest/analysis/running-pipelines/cr-flex-multi-frp). Reference genome used for mapping was hg38 genome and gene annotation was performed on Ensembl 98. All subsequent data processing was conducted in R using Bioconductor (R version 4.2.2, Bioconductor version 3.16).

### Alignment and demultiplexing of single-cell 3’ gene expression data

Sequencing data were mapped to the hg38 genome using the STARsolo framework (v2.7.10a) and gene annotation was performed with a custom GTF file (Ensembl 104), filtered to include only transcripts of biotypes “protein_coding,” “lncRNA,” and all B- and T-cell receptor gene transcripts. The cell barcode whitelist for STAR was provided by 10x (file: 737K-august-2016.txt). Non-standard command line options for STAR included parameters for filtering, mapping, output formatting, and handling cell barcodes and UMIs, such as “– outFilteType BySJout,” “–outFilterMultimapNmax 10,” “–outSAMtype BAM SortedByCoordinate,” and various options specific to the handling of cell barcodes (CB) and unique molecular identifiers (UMIs). All subsequent data processing was conducted in R using Bioconductor (R version 4.2.2, Bioconductor version 3.16).

### Quality controls and normalization of single-cell fixed RNA-Profiling data

For each cell, three quality metrics were calculated: percentage of mitochondrial genes, total number of reads, and total number of genes using scuttle (v1.8.4). Cells with fewer than 2,000 total reads, fewer than 1,000 detected genes, more than 10% of mitochondrial genes, or identified as doublets by the scDblFinder (v1.12.0) were excluded from further analysis. Data normalization was then performed sample-wise using the R package scran (v1.26.2). In order to calculate pool-based normalization factors, cells were first split into sensible clusters using the quickCluster() function with default parameters and cluster-specific size factors were calculated with the *computeSumFactors()* function. Data were then normalized with *logNormCounts()* using previously computed cluster-specific size factors.

### Quality controls and normalization of single-cell 3’ Gene Expression data

Empty droplets were filtered out using the emptyDrops(niters = 5000) function from the DropletUtils package (v1.18.1). Quality control was then conducted similarly to the single-cell fixed RNA profiling data. Cells with fewer than 10,000 total reads, fewer than 1,000 detected genes, more than 25% mitochondrial content, or those identified as doublets by scDblFinder were excluded from further analysis.

### Analysis of public scRNA-seq data

When made available by their authors, filtered count matrix which already underwent quality controls steps were used as input for subsequent steps. When such objects were not directly available, raw count matrices were used as in input to reproduce quality controls as close as described in the original publication. All datasets were first separated by patient-ID and normalized using the same method as for in-house data to ensure consistency.

PCa-derived organoid data from Huang *et al.* ^24^ were accessed via the GitHub page of the lab as a pre-filtered Seurat object (https://github.com/franklinhuanglab/Clubcell_Organoid). The filtered count matrix was extracted from the Seurat object and converted into a SingleCellExperiment (SCE) object and normalization was performed. Post-normalization, only cells cultured with DHT (1 nM) were retained, while cells subjected to other treatments were excluded for subsequent integration analyses.

PCa and healthy-derived organoid data from Song *et al.* ^26^ were accessed via GEO online portal (<u>GSM5353249</u> to <u>GSM5353266</u>) as raw count matrices. Quality control filtering was applied based on the following criteria: cells with less than 500 reads, less than 300 detected genes, or more than 20% mitochondrial content were excluded. The filtered data were then converted to a SCE object, separted by patient-ID and normalization was performed.

Prostate-derived organoid data from McCray *et al.* ^39^ were accessed via GEO online portal (<u>GSM3735994</u> ) as a raw count matrix. Quality control filtering was applied based on the following criteria: cells with less than 11,000 reads, less than 500 detected genes, and more than 20% mitochondrial content. The filtered data were then converted to a SCE object, separated by patient-ID and normalization was performed.

PCa tissue data from Chen *et al.* ^25^ were accessed via GEO online portal (<u>GSM4203181</u>) as a raw count matrix. Quality control filtering was applied based on the following criteria: cells with less than 200 or more than 5,218 detected genes, more than 20% mitochondrial content and fewer than 56 housekeeping genes. The filtered data were then converted to a SCE object, separated by patient-ID and normalization was performed.

Healthy prostate tissue data from Henry *et al*. ^22^ were accessed via the UCSC Cell Browser (https://cells.ucsc.edu/?ds=prostate-prostatic-urethra). The count matrix (exprMatrix.tsv.gz) and cell type annotation (meta.tsv) were downloaded from the “Info & Download” subpage. After creating a SCE object, normalization was performed.

Healthy prostate tissue data from the *Tabula Sapiens* study ^38^ were accessed via the CELLxGENE online portal (https://cellxgene.cziscience.com/collections) as a filtered Seurat object. After converting it to a SCE object, normalization was performed.

### Multiple sample integration

To integrate multiple samples, the per-sample normalized objects were first harmonized by retaining only genes shared across all SCE objects. Next, to account for differences in sequencing depth between batches, all SCE objects were rescaled using the MultiBatchNorm(min.mean = 0.1) function from the Batchelor R package (v1.14.1). Using scran (v1.26.2), variance modelling of gene expression was performed for each sample individually with the modelGeneVar() function, results were merged with combineVar(), and the top 1000 most highly variable genes were selected getTopHVGs() for subsequent analysis. The function fastMNN(k = 300, d = 20) built within Batchelor was used to apply batch correction. In order to obtain a two-dimensional representation of the integrated data, UMAP was then performed using the 20 first principal components of the batch corrected data using RunUMAP(n_neighbors = 300, min_dist = 0.5).

### Clustering analysis

To identify transcriptomically-distinct subpopulations within our cohort of Matrigel- and ECMf- derived organoids, clustering was performed on the integrated data using the clusterCells() function from the scran R package (v1.26.2). The clustering was conducted with the Walktrap algorithm, exploring different values of the k parameter ranging from 10 to 50 in increments of 10. The optimal clustering resolution was achieved with k = 30, which resulted in 8 distinct clusters. This number of clusters was selected to approximate the known diversity of epithelial subtypes in the prostate and PCa, which can range from 5 to 10, including variations observed among cells at different stages of the cell division cycle ^22^.

### Differential gene expression analysis

To identify differentially expressed (DE) genes across cell groups, the FindAllMarkers() function from the Seurat package (v5.0.1) was employed using the Wilcoxon test. Various parameters were adjusted, including the minimum number of cells expressing a gene and the log fold change threshold depending on groups compared. The results were visualized as heatmaps using the ComplexHeatmap package (v2.14.0), with genes sorted by ascending adjusted p-value.

### Signature score calculation

After clustering analysis, clusters were characterized by identifying enrichment for particular gene signatures. Gene signatures were retrieved from the MSigDB database with the help of the R package msigdbr (v7.5.1) and the function AddModuleScore() built within Seurat (v5.0.1) was then used to generate a signature score for each cell. The distribution of these scores was evaluated across clusters and a one-way ANOVA test was conducted to assess differences in signature enrichment. This was followed by post-hoc t-tests, using the cluster with the lowest mean signature score as the reference group.

### Cell-type annotation

Based on Expression data, DEA, Signature scores and Pseudotime analysis clusters 1,8 were manually assigned as “Tumor”, clusters 2,5,6,7 as “Basal/Hillock”, cluster 3 as “Transitioning” and cluster 4 as “Club”. Cell-type annotation for parental tissues and cells from previously published datasets was performed using the SingleR() function of the SingleR package (v2.0.0), using SCE object containing cell-type annotations for reference and “de.method = wilcox ”.

### Copy number variation analysis

Copy number variation (CNV) analysis was performed using the inferCNV R package (v0.8.2). The infercnvObject was constructed using normal prostate-derived organoids from the publications by Song et al. and McCray et al. ^26,39^, with mitochondrial chromosomes excluded from the analysis. The run() function was executed with the following non-default parameters: “cutoff = 0.1”, “analysis_mode = samples”, “min_cells_per_gene = 20”, “scale_data = TRUE”, and “HMM = TRUE”. A score was then calculated for each cell to represent the proportion of genes whose expression values fall outside the range of 0.90 to 1.10, which may indicate potential CNVs in the genome of these cells.

### Pseudotime analysis

Integrated single-cell RNA sequencing (scRNA-seq) data from Matrigel- and ECMf-derived organoids were used for pseudotime analysis. Pseudotime trajectory inference and lineage analysis were performed using the slingshot (v2.6.0) using cluster labels as inputs. The getLineages() function was initialized from cluster “6,” and getCurves(approx_points = 300, thresh = 0.01, stretch = 0.8) was used to fit principal curves, capturing developmental trajectories. To explore gene expression dynamics along pseudotime, a series of generalized additive models (GAMs) was fitted using the fitGAM() function from the tradeSeq package (v1.8.0). Finally, differential expression between trajectory start and end points was assessed with startVsEndTest() with default parameters to identify key progenitor markers.

### Statistical analysis and data representation

Besides bioinformatic analyses described above, statistical analyses were conducted using a paired t-test (Fig. 1) or a Wilcoxon test (Fig. 3i). P values lesser than 0.05 were considered significant. The Prism (version 10.1.1, 270), and R-studio software were used for statistical analyses and data representation.

### Data availability

The raw FASTQ files single-cell RNA sequencing generated in this study have been submitted to the European Genome-phenome Archive. Other datasets and their accessibility are described in details in the paragraphs above. Spatial transcriptomic data from Erickson et. al ^35^ were re-analyzed and visual representation of gene expression was generated using the Loupe Browser 7 (version 7.0.1).

## Acknowledgments

We thank all patients who consented to participate to the study and all clinicians, pathologists, and study nurses involved in the samples acquisition. We acknowledge the IHC/FISH facility of the Institute of Pathology (Petra Hirschmann, Bruno Grili), as well as the microscopy (Ewelina Bartoszek, Loic Sauteur) and bioinformatic facilities (Julien Roux, Florian Geier) of the DBM for technical and scientific support. We thank the Next Generation Sequencing Platform of the University of Bern for performing the scRNA sequencing. We are grateful to Yu Chen (MSKCC, US) for kindly providing the MSK-PCa2 organoid line, to Dimitrios Doultsinos (University of Oxford, UK) for support with the spatial RNA-seq data, and to Gaëlle Schappler and Renaud Mével for experimental support. We thank Wouter Karthaus (EPFL/ISREC, CH) for critical review of the manuscript. Financial support was provided by funding from Krebsliga beider Basel (KLbB-5329-03-2021 to CL), the Science National Science Foundation (320030_205086 to CL), the Swiss Cancer Research Foundation (KFS-5552-02-2022 to AM, CL, CAR), and the Department of Surgery of the University Hospital Basel (*PMC* Platform to CL). Components of several figures were created with BioRender.com.

## Author contributions

R.D., C.L., and R.P. conceived the study. C.L. supervised the study. R.D., R.P., J.W., R.S. designed and conducted experiments. R.D. and R.P. analyzed data and performed data visualization. A.J.T., T.Z., K.D.M., S.S., T.V., H.S., A.M., C.A.R., L.B. enrolled patients in the study and all contributed to patient samples collection and clinical data. T.V. and L.B. performed the histopathological analyses. A.D.L. provided technical support and spatial data. R.D., C.L., and R.P. wrote the initial manuscript and all the authors contributed to the final version.

## Competing interests

All authors declare no competing interest.

## Notes

### Competing Interest Statement

The authors have declared no competing interest.

