## Supplementary Material for "ECM-free patient-derived organoids preserve diverse prostate cancer lineages and uncover *in vitro*-enriched cell types"

#### Supplementary Figures

Figure S1. Modulating medium factors and the extracellular matrix affects the cellular composition and viability of PDOs

Figure S2. Phenotypic analysis of PDOs grown in ECM-free or Matrigel conditions

Figure S3. Cluster-specific features

Figure S4. PCa-associated and androgen response signatures in cell clusters

Figure S5. Expression of selected genes and lineage trajectory in identified cell clusters

Figure S6. Expression of specific marker genes in *in vitro*-associated cell clusters and in patient samples *in situ*

Figure S7. Patient-specific and cluster-associated transcriptomic features

#### Supplementary Tables

Table S1. Patients and samples characteristics (excel file)

Table S2. Quantification results for Figure 1e - f

Table S3. Quantification results for Figure 1g

Table S4. List of antibodies used in this study

#### Supplementary Dataset

Supplementary dataset 1. Differential expression analysis (excel file)

#### Supplementary Figures

### Fig. S1

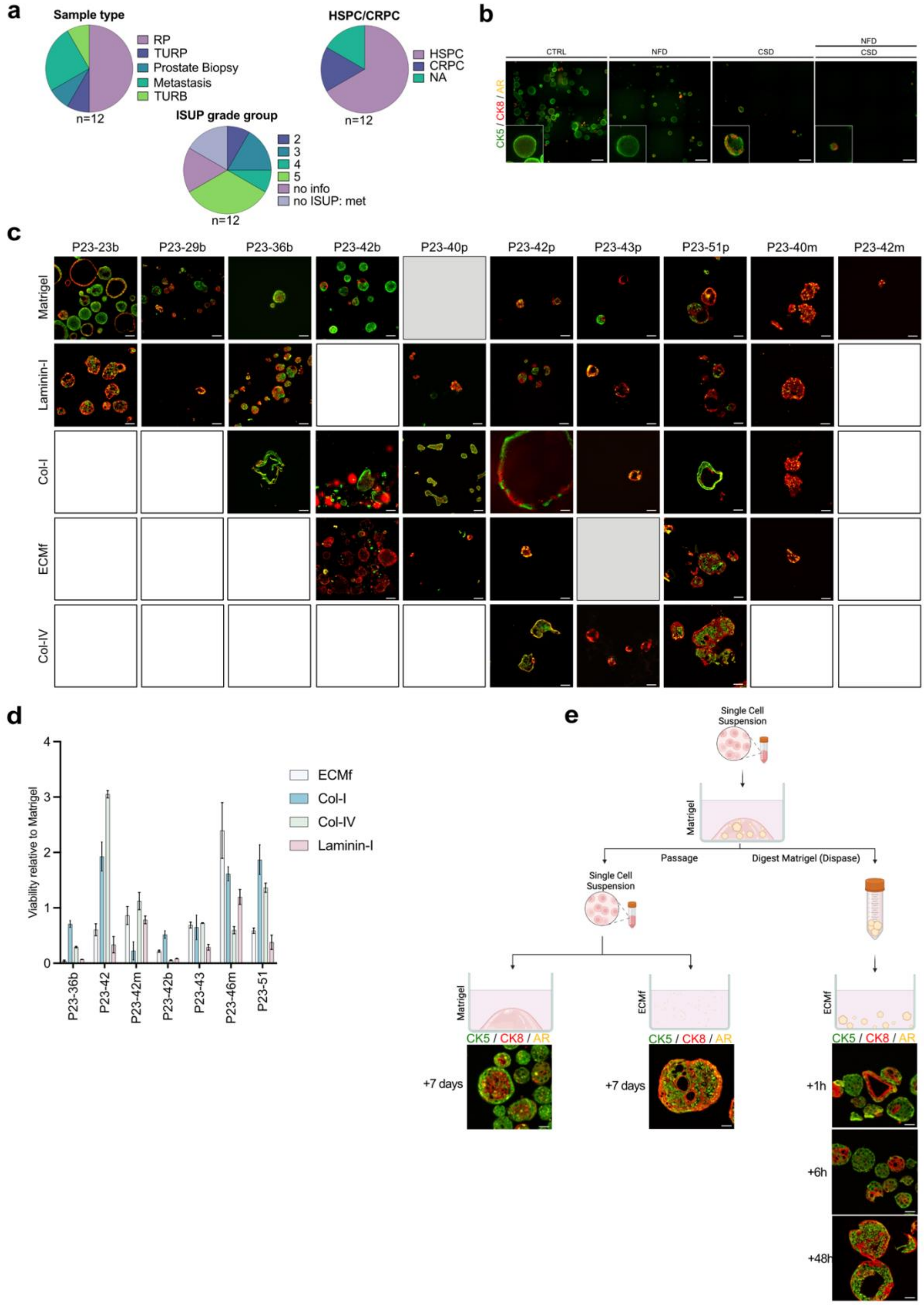

**Figure S1. Modulating medium factors and the extracellular matrix affects the cellular composition and viability of PDOs. a** PCa patient sample cohort (n=12) used to generate PDOs classified as "Short-term growth: confirmed cancer features" in Figure 1a. **b-c** Whole-mount immunofluorescence images (b) or FFPE sections immunofluorescence images (c) of PDOS cultured in different conditions. CTRL: Published control conditions, NFD: Niche factor deprivation, CSD: Carbon source deprivation. Organoids were stained with antibodies recognizing cytokeratin 5 (CK5), cytokeratin 8 (CK8), and androgen receptor (AR). Insets show higher magnification of a representative area. Sample types labelled as follows: b: benign-like PDOs, p: Primary PCa, m: PCa metastasis. Scale bars represent 50 μm. White squares indicate no growth, and grey squares indicate no additional data. **d** Viability analysis of organoids cultures in different matrix conditions. Data are represented as means relative to the Matrigel control and the error bars represent standard deviations (SD) for three technical replicates. b = benign organoid lines. m = metastatic specimen. **e** Top: Experimental workflow of pre-formed organoid polarity reversal experiment. Bottom: Immunofluorescence images of P23-23 PDOs stained with antibodies recognizing cytokeratin 5 (CK5), cytokeratin 8 (CK8), and androgen receptor (AR). Scale bars represent 50 μm.

#### Fig. S2

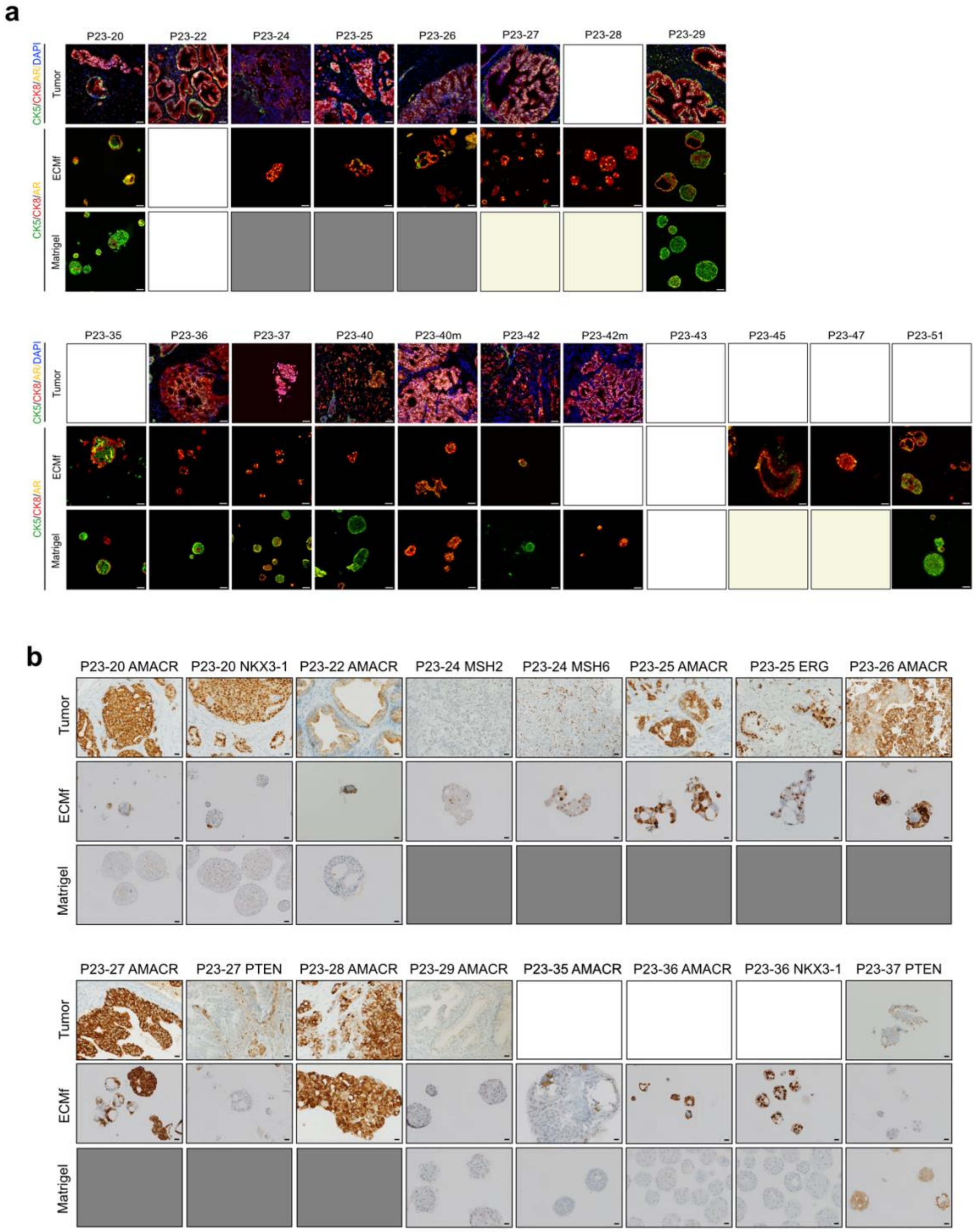

**Figure S2. Phenotypic analysis of PDOs grown in ECM-free or Matrigel conditions. a** Immunofluorescence images of patient tumor tissues and matched PDOS cultured in either condition. Organoids were stained with antibodies recognizing cytokeratin 5 (CK5), cytokeratin

8 (CK8), and androgen receptor (AR). Insets show higher magnification of a representative area. Scale bars represent 50 μm. White square: no additional image. Grey square: no growth. Yellow square: Not enough material to test both conditions. **b** IHC analysis for the indicated antibodies in patient tumor tissues and matched PDOS cultured in either condition. Scale bars represent 20 μm.

### Fig. S3

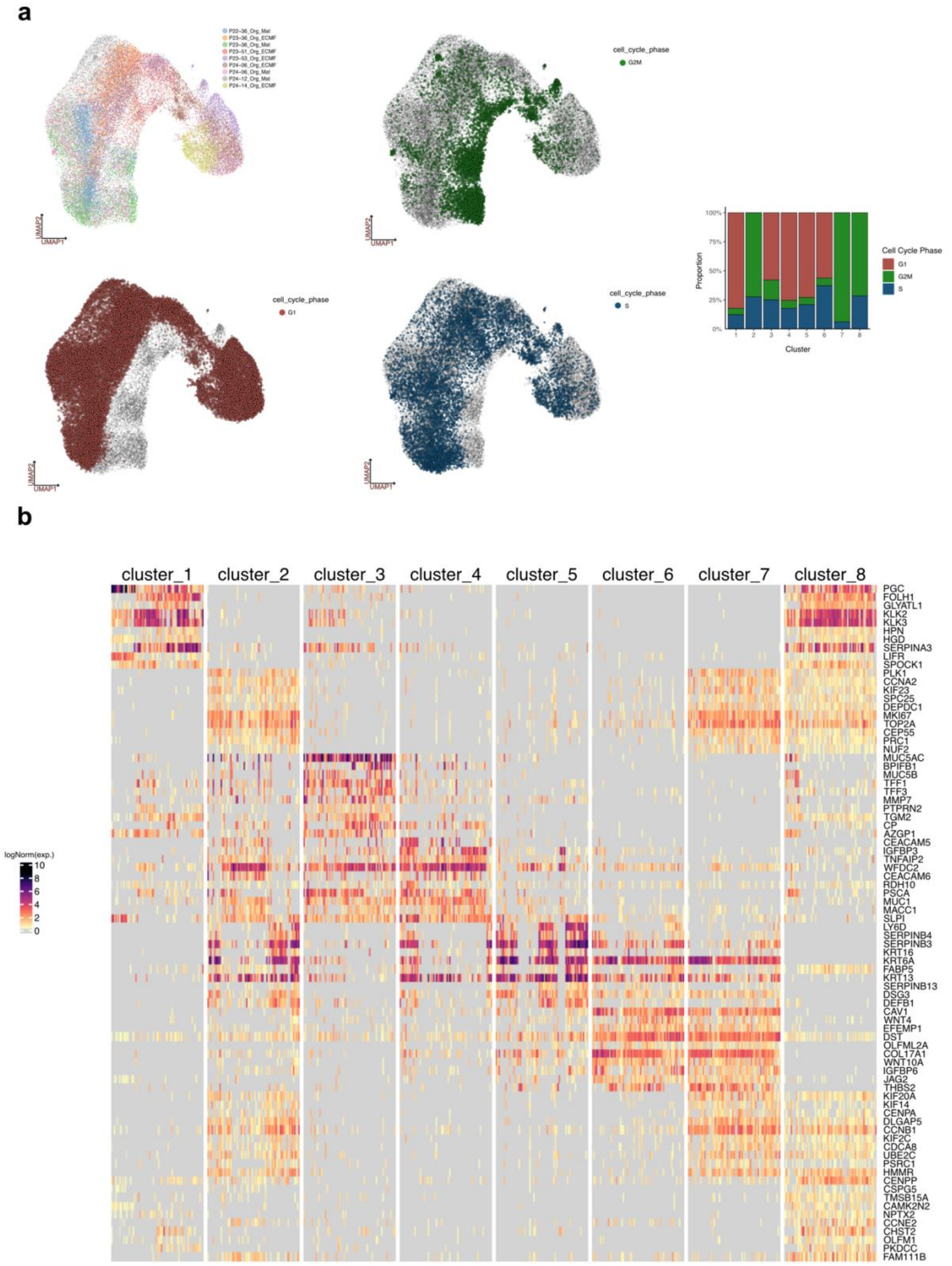

**Figure S3. Cluster-specific features. a** (left) UMAP projection of organoid-derived cells color-coded by sample identity or by cell cycle phase. (right) Cell cycle phase proportion in each identified cluster. Cell counts in each cluster are normalized to 100%. **b** Top 10 differentially-expressed genes between clusters represented as a single cell heatmap plot.

#### Fig. S4

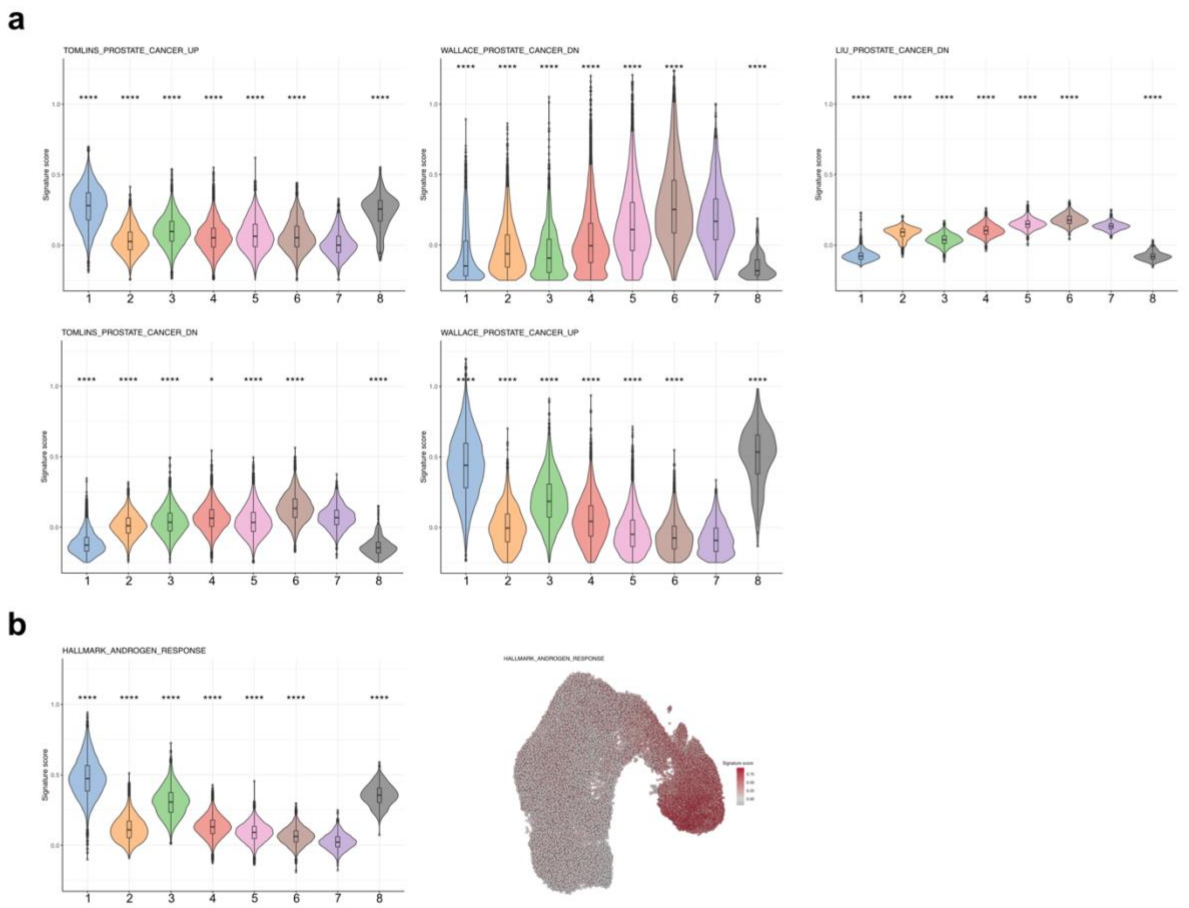

**Figure S4. PCa-associated and androgen response signatures in cell clusters. a** Signature scores per cluster, represented as violin plots for the eight clusters. From left to right: “Tomlins prostate cancer up”, (Collection C2: Curated, gsea_msigdb), “Wallace prostate cancer DN”, (Collection C2: Curated, gsea_msigdb), “Liu prostate cancer DN”, (Collection C2: Curated, gsea_msigdb), “Tomlins prostate cancer DN”, (Collection C2: Curated, gsea_msigdb), “Wallace prostate cancer up”, (Collection C2: Curated, gsea_msigdb). Anova (p<2.2e-16) followed by t-tests were performed. ns: P>0.05, *:P<0.05, **:P<0.01, ***:P<0.001, ****: P<0.0001. **b** (Left) “Hallmark androgen Response” signature score per cluster, represented as violin plots for the eight clusters (Collection H: Hallmark, gsea_msigdb). (Right) UMAP representation color-coded by intensity of signature score.

### Fig. S5

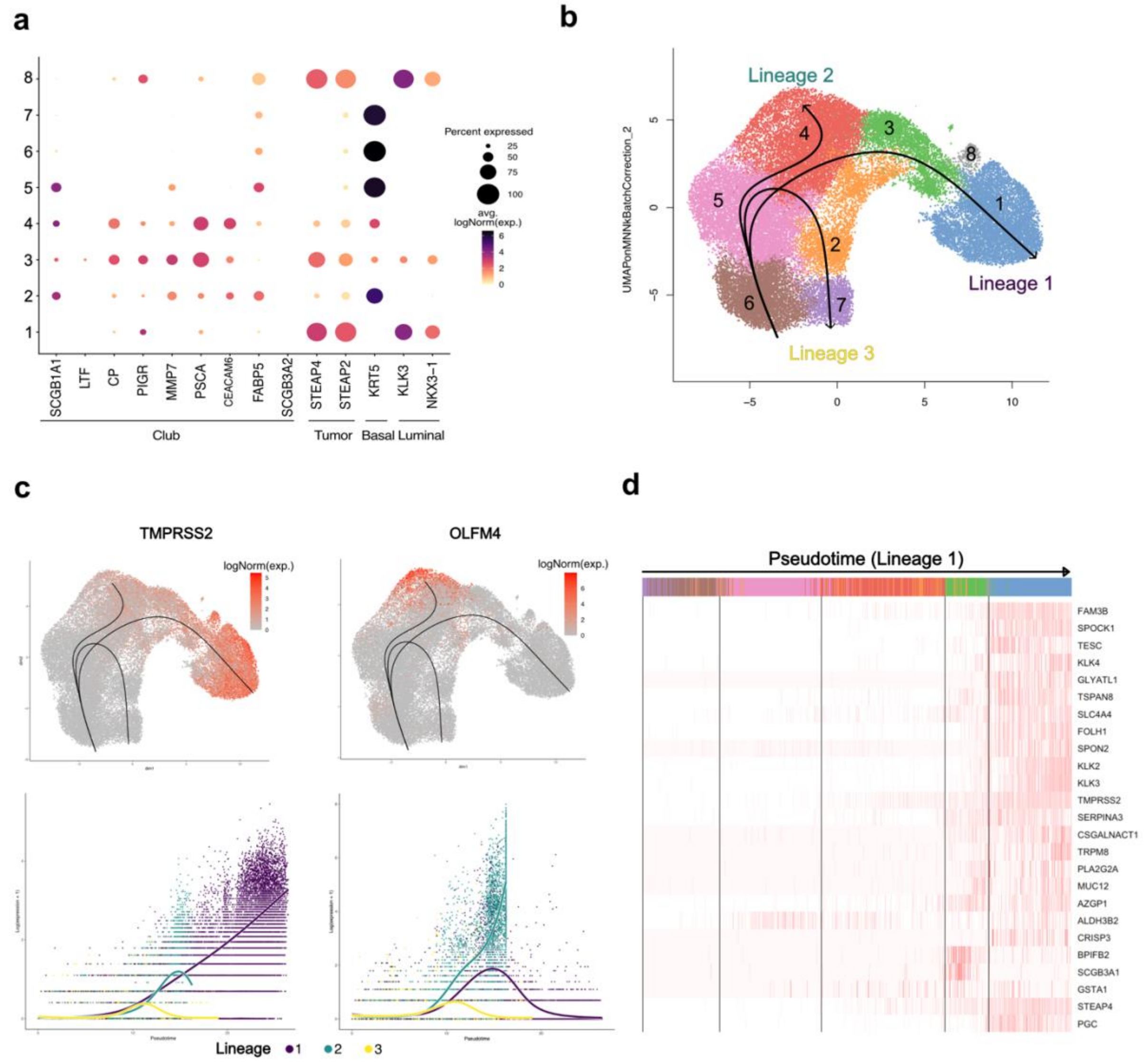

**Figure S5. Expression of selected genes and lineage trajectory in identified cell clusters. a** Dotplot representing expression of selected genes in each cluster. The color represents scaled average expression of marker genes in each cell type and the size indicates the proportion of cells expressing each gene. **b** Pseudotime trajectory lineages determined with cluster 6 as starting point. **c** (Top) UMAP projection of organoid-derived cells color-coded by logged count expression of selected genes. (Bottom) Expression of selected genes across lineage trajectories. **d** Most differentially-expressed genes across lineage 1 represented as a heatmap.

#### Fig. S6

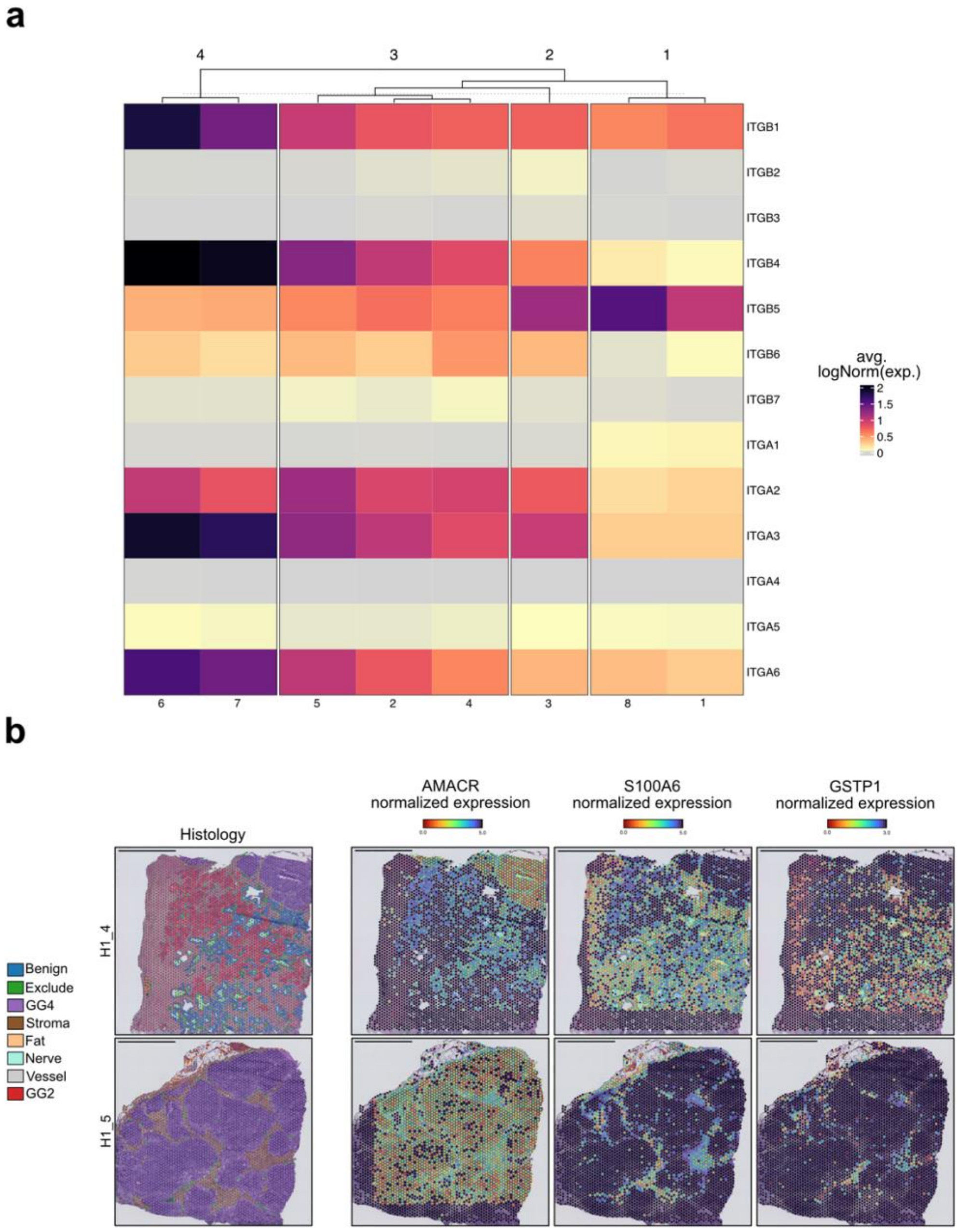

**Figure S6. Expression of specific marker genes in *in vitro*-associated cell clusters and in patient samples *in situ*. a** Heatmap plot representing unsupervised hierarchical clustering analysis of selected integrin genes expression in the distinct *in vitro* clusters. **b** Spatial transcriptomics (Visium ST) data from H1_5 and H1_4 prostatectomy sections, with spot-level pathological classification of areas (left), and log-normalized expression of AMACR, S100A6 and GSTP1 (right). Scale bars represent 2.5 mm. Data from Erickson et al., *Nature 2022*.

### Fig. S7

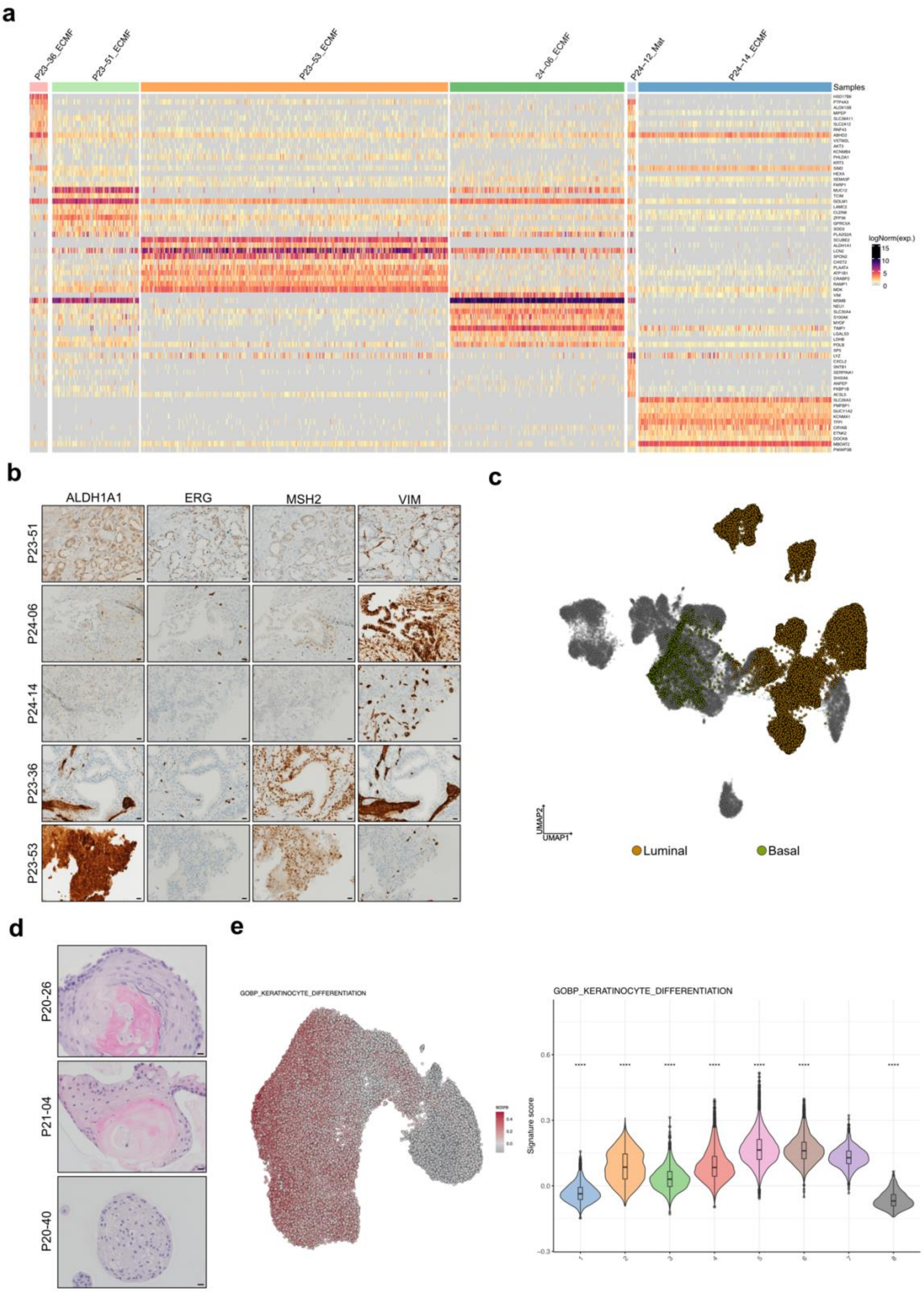
a
P23-36_ECMF
P23-51_ECMF
P23-53_ECMF
24-06_ECMF
P24-12_Mat
P24-14_ECMF
Samples
logNorm(exp.)
15
10
5
0
b
ALDH1A1
ERG
MSH2
VIM
P23-51
P24-06
P24-14
P23-36
P23-53
c
UMAP2
UMAP1
Luminal
Basal
d
P20-26
P21-04
P20-40
e
GOBP_KERATINOCYTE_DIFFERENTIATION
GOBP_KERATINOCYTE_DIFFERENTIATION
Signature score
0.6
0.3
0.0
-0.3
****
1
2
3
4
5
6
7
8

**Figure S7. Patient-specific and cluster-associated transcriptomic features. a** Top 10 differentially-expressed genes in clusters 1,8 (tumor) of each patient-specific ECMf organoid culture represented as a single cell heatmap plot. **b** IHC analysis for the indicated antibodies in original patient samples. Scale bars represent 20 μm. **c** UMAP projection of PCa-derived cells of the Chen dataset color-coded by cell type (orange or yellow) as assigned in their original population. **d** H&E staining of organoids derived from three PCa tissues and cultured in Matrigel. Scale bars represent 20 μm. **e** (Left) UMAP representation color-coded by intensity of the "GOBP Keratinocyte Differentiation" (Collection C5: Ontology, msigdb) signature score. (Right) signature score per cluster, represented as violin plots for the eight clusters.

| | Table S2: Quantification results for Figure 1e - f | | | | |
|---|---|---|---|---|---|
| | | Luminal | Basal | Mixed | Total |
| CTRL | P23-20 | *5* | *19* | *7* | 31 |
| | P23-21 | *5* | *116* | *12* | 133 |
| | P23-22 | *3* | *25* | *9* | 37 |
| NFD | P23-20 | *2* | *6* | *5l* | 13 |
| | P23-21 | *5* | *31* | *13* | 49 |
| | P23-22 | *3* | *17* | *7* | 27 |
| CSD | P23-20 | *2* | *1* | *2* | 5 |
| | P23-21 | *2* | *7* | *2* | 11 |
| | P23-22 | *0* | *7* | *0* | 7 |
| NFD + CSD | P23-20 | *2* | *1* | *0* | 3 |
| | P23-21 | *2* | *0* | *0* | 2 |
| | P23-22 | *0* | *0* | *0* | 0 |

| | Table S3: Quantification results for Figure 1g | | | | |
|---|---|---|---|---|---|
| | | Luminal | Basal | Mixed | Total |
| Matrigel | P23-23b | 2 | 14 | 4 | *20* |
| | P23-29b | 8 | 5 | 11 | *25* |
| | P23-36b | 0 | 90 | 10 | 100 |
| | P23-40 | 3 | 23 | 4 | 30 |
| | P23-40m | 8 | 0 | 0 | 8 |
| | P23-42 | 11 | 16 | 7 | *34* |
| | P23-42m | 9 | 0 | 0 | 9 |
| | P23-42b | 25 | 96 | 113 | 234 |
| | P23-43 | 5 | 0 | 2 | *7* |
| | P23-51 | 19 | 23 | 7 | *49* |
| Laminin-I | P23-23b | 0 | 0 | 9 | *9* |
| | P23-29b | 3 | 0 | 0 | *3* |
| | P23-36b | 6 | 0 | 18 | 24 |
| | P23-40 | 6 | 0 | 0 | 6 |
| | P23-40m | 3 | 0 | 0 | 3 |
| | P23-42 | 4 | 0 | 3 | *7* |
| | P23-42m | 0 | 0 | 0 | 0 |
| | P23-42b | 0 | 0 | 0 | 0 |
| | P23-43 | 27 | 0 | 2 | *29* |
| | P53-51 | 4 | 0 | 2 | *6* |
| Col-I | P23-23b | 0 | 0 | 0 | 0 |
| | P23-29b | 0 | 0 | 0 | 0 |
| | P23-36b | 0 | 9 | 1 | 1 |
| | P23-40 | 0 | 15 | 6 | 21 |
| | P23-40m | 5 | 0 | 0 | 5 |
| | P23-42 | 0 | 1 | 0 | *1* |
| | P23-42m | 0 | 0 | 0 | 0 |
| | P23-42b | 0 | 1 | 0 | 1 |
| | P23-43 | 0 | 1 | 0 | *1* |
| | P23-51 | 0 | 3 | 3 | *6* |
| Col-IV | P23-23b | 0 | 0 | 0 | 0 |
| | P23-29b | 0 | 0 | 0 | 0 |
| | P23-36b | 0 | 0 | 0 | 0 |
| | P23-40 | 0 | 0 | 0 | 0 |
| | P23-40m | 0 | 0 | 0 | 0 |
| | P23-42 | 2 | 2 | 11 | *15* |
| | P23-42m | 0 | 0 | 0 | 0 |
| | P23-42b | 0 | 0 | 0 | 0 |

| | | | | | |
|---|---|---|---|---|---|
| | P23-43 | 1 | 0 | 9 | *10* |
| | P23-51 | 0 | 0 | 13 | *13* |
| ECMf | P23-23b | 0 | 0 | 0 | 0 |
| | P23-29b | 0 | 0 | 0 | 0 |
| | P23-36b | 0 | 0 | 0 | 0 |
| | P23-40 | 5 | 0 | 0 | 5 |
| | P23-40m | 5 | 0 | 0 | 5 |
| | P23-42 | 18 | 0 | 0 | *18* |
| | P23-42m | 0 | 0 | 0 | 0 |
| | P23-42b | 9 | 0 | 0 | 9 |
| | P23-43 | 9 | 0 | 0 | *9* |
| | P23-51 | 12 | 0 | 14 | *26* |

| Table S4: List of antibodies used in this study | | | | |
|---|---|---|---|---|
| **Primary antibodies used for Immunohistochemistry** | | | | |
| Antigen | Type | Supplier* | Catalog # | Dilution |
| ERG | Rabbit mAb | Ventana | 790-4576 | ready to use |
| PTEN | Rabbit mAb | Ventana | 790-5097 | ready to use |
| NKX3.1 | Rabbit mAb | Ventana | 760-5086 | ready to use |
| AMACR | Rabbit mAb | Ventana | 790-6011 | ready to use |
| GSTP1 | Mouse mAb | Leica | NCL-GSTpi-438 | ready to use |
| MSH2 | Mouse mAb | Ventana | 760-5093 | ready to use |
| MSH6 | Mouse mAb | Ventana | 760-4389 | ready to use |
| Vimentin | Mouse mAb | Ventana | 790-2917 | ready to use |
| ALDH1A1 | Rabbit mAb | Abcam | Ab52492 | 1:50 |
| **Primary antibodies used for Immunofluorescence** | | | | |
| Antigen | Type | Supplier* | Catalog # | Dilution |
| Androgen Receptor | Rabbit mAb | Abcam | ab133273 | 1:200 |
| Cytokeratin 5 | Chicken pAb | Biolegend | 905901 | 1:1000 |
| Cytokeratin 8 | Mouse mAb | Biolegend | 904801 | 1:1000 |
| **Secondary antibodies for immunofluorescence** | | | | |
| Antigen | Fluorochrome | Company | Catalog # | Dilution |
| Goat anti-Chicken IgY | Alexa Fluor® 555 | Invitrogen | A-21437 | 1:1000 |
| Goat anti-Chicken IgY | Alexa Fluor® 488 | Invitrogen | A-11039 | 1:1000 |
| Goat anti-Rabbit IgG | Alexa Fluor® 555 | Invitrogen | A-21428 | 1:1000 |
| Goat anti-Mouse IgG | Alexa Fluor® 647 | Invitrogen | A-21237 | 1:1000 |
